# Novel antibiotic mode of action by repression of promoter isomerisation

**DOI:** 10.1101/2020.12.31.424950

**Authors:** Leena Kerr, Douglas F. Browning, Kimon Lemonidis, Talal Salih, Iain S. Hunter, Colin J. Suckling, Nicholas P. Tucker

## Abstract

Rising levels of antibiotic resistance dictate that new antibiotics with novel modes of action must be found. Here, we investigated the mode of action of a novel antibiotic that is a member of a family of synthetic DNA minor groove binding (MGB) molecules. MGB-BP-3 has successfully completed a Phase II clinical trial in humans as an orally administered drug for the treatment of chronic *Clostridioides* (*Clostridium) difficile* infections, where it outperformed the existing benchmark (vancomycin). MGB-BP-3 is active against a variety of Gram-positive pathogens including *Staphylococcus aureus,* which was used as the model for this study. The transcriptomic response of *S. aureus* to MGB-BP-3 identified downregulated promoters. DNase I and permanganate footprinting demonstrated binding to essential SigA promoters and the inhibition of promoter isomerisation by RNA polymerase holoenzyme. Promoters controlling DNA replication and peptidoglycan biosynthesis are amongst those affected by MGB-BP-3. Thus, MGB-BP-3 binds to and inhibits multiple essential promoters on the *S. aureus* chromosome, suggesting that evolution of resistance by drug target mutation should be unlikely. In confirmation, laboratory-directed evolution against sub-inhibitory concentrations of MGB-BP-3 resulted in no resistance whereas resistance to the single target RNA-polymerase inhibitor rifampicin arose rapidly.

## Introduction

DNA minor-groove binding (MGB) drugs have a variety of effects on infectious agents including bacteria, fungi, parasites and viruses [1]. Distamycin, a natural product made by *Streptomyces*, has been used as a chemical biology probe to examine the structure of AT-rich promoter UP elements in *Escherichia coli* and has shown promise as an anti-cancer agent [2, 3]. Distamycin inhibits the binding of the transcriptional machinery to DNA [4, 5], but its toxicity to humans has prevented it from being developed as an anti-infective agent. Using distamycin as a design concept, a family of synthetic MGBs was synthesised [*e.g.* 6]. These structural variants retained DNA-binding and anti-Gram-positive activity but lacked human toxicity [6]. One of these, MGB-BP-3 (Figure 1a), has been taken forward for clinical development. It has strong (<1 μg/ml) antibacterial activity against methicillin-resistant and -susceptible *Staphylococcus* species, pathogenic *Streptococcus* species, vancomycin-resistant and susceptible *Enterococcus,* and *Clostridioides* (*Clostridium*) *difficile*. Its oral formulation, developed for the treatment of *C. difficile* infections, has successfully completed a Phase II clinical trial. Subsequently, different compounds have been found to be active against a wide variety of pathogens and some cancer cell lines [7–11]. Antibacterial activity of MGB-BP-3 is confined to Gram-positive bacteria and dose response curves show a steep decrease in viability over a narrow concentration range, suggesting a catastrophic failure in the bacterium rather than an interaction with a single receptor typified by a sigmoidal dose-response curve. Members of this MGB family have typically been shown to bind to 6–8 base pairs in dsDNA [6, 12]. Whilst it can be reasonably anticipated that the main mode of action of MGB-BP-3 is by binding to DNA, the detailed biological consequences of binding to DNA that eventually lead to cell death cannot be predicted and need further investigation. Moreover, a short minor groove binder such as MGB-BP-3 could bind to many sites on the bacterial genome with the potential to interfere with a number of essential biological processes. Here, we have used RNA-Sequencing to probe the transcriptional consequences of MGB-BP-3 binding to the *S. aureus* chromosome. This approach, along with qPCR melt analysis, phenotypic microarrays, DNase I footprinting and potassium permanganate footprinting allows the identification of promoters that are sensitive to MGB-BP-3 shedding new light on its mode of action against *S. aureus*.

**Figure 1:**
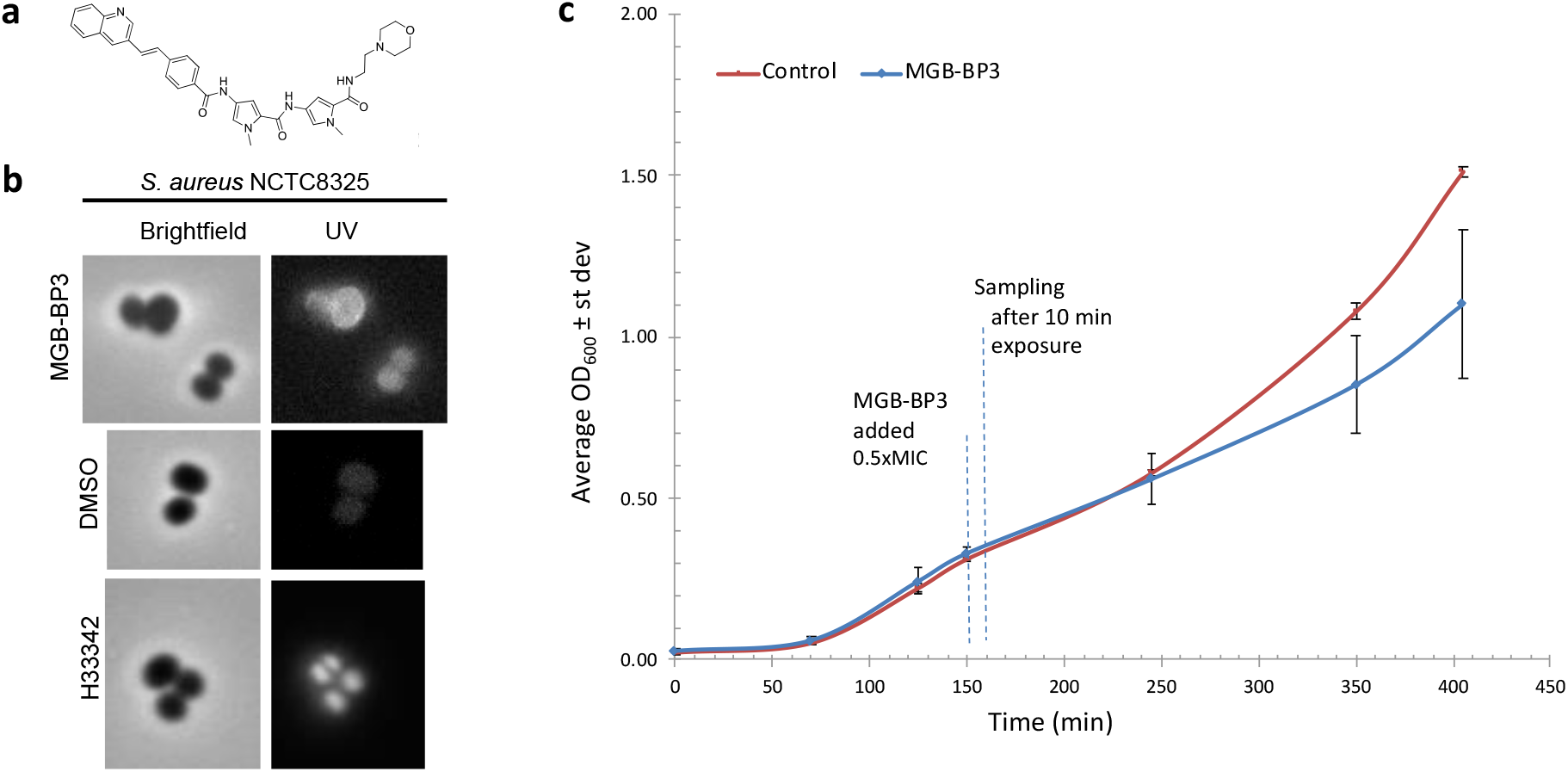
Sub-lethal exposure of *S. aureus* to MGB-BP-3. **a**. Structure of MGB-BP-3. **b**. Microscopy images showing binding of MGB-BP-3 to *S. aureus* cells under UV filter set. DNA binding dye, H33342, used as a positive control and DMSO as a negative control. **c**. *S. aureus* NCTC8325 growth, as measured by OD_600_ MGB-BP-3 (0.5 x MIC) treated and untreated (control) cultures. Cells were harvested for RNA-Seq experiment 10 min after exposure. Error bars, s.d., *n* = 3

These data indicate multiple chromosomal binding sites, suggesting that generation of resistance to this type of drug would be significantly less than for drugs with a single target. In order to test this hypothesis we used a directed evolution approach to compare the evolvability of resistance to MGB-BP-3 and rifampicin.

Together, these results provide a comprehensive profile of the effect of MGB-BP-3 on *S. aureus* in culture and support the concept of multiple antibacterial actions via selective, but simultaneous, inhibition of a subset of promoters.

## Materials and Methods

### Bacterial strain and growth conditions

*S. aureus* subsp. *aureus* strain NCTC8325 from the HPA Culture Collection was used throughout this study with the exception of the resistance/evolution experiment for which *S. aureus* ATCC 43300 (resistant to methicillin and oxacillin) was used. Bacteria were grown for transcriptomic analyses according to the following: pre-cultures were prepared by inoculation from a frozen bead stock (Microbank, Fisher Scientific UK) to 5 ml of Tryptic Soya Broth (Sigma-Aldrich) and incubated overnight. Six 250 ml shake flasks with 50 ml of cation-adjusted Mueller-Hinton broth 2 (Sigma-Aldrich) were inoculated to OD_600_ of 0.05 using a pre-culture. All cultures were incubated at 250 rpm and 37°C. At OD_600_ of 0.3, 0.5 x MIC of MGB-BP-3 dissolved in DMSO was added to the cultures in triplicate (treated samples), whereas control samples (also in triplicate) where treated with same volume of DMSO only (untreated samples). Samples of 10 ml were withdrawn 10 min after addition of antibiotic. The samples were immediately transferred to RNAprotect Bacteria Reagent (Qiagen) following the supplier’s instructions. The MIC was determined using the broth dilution method on 96-well plate in cation-adjusted Mueller-Hinton broth 2 (Sigma-Aldrich).

### Microscopy

*S. aureus* NCTC8325 cells, grown to 0.3-0.4 OD_600_ in nutrient broth (Oxoid), were treated with MGB-BP-3 (3 μg/ml) for *ca.* 40 mins. DNA binding dye Hoechst 33342 was used as a positive control (at 14 μg/ml) and DMSO was used as a negative control. Brightfield and fluorescent images (UV filter set) were captured using a Nikon TE2000S inverted fluorescence microscopy with IPLab scientific imaging software version 3.7 (Scanalytics, Inc., Rockville, USA).

### RNA extraction and library preparation

Total RNA was extracted using a bacterial RiboPure RNA Purification Kit (AM1925, ThermoFisher Scientific) immediately after samples were collected. In brief, the cells were disrupted mechanically using Zirconia beads, total RNA was then extracted in phenol and purified using glass-fibre filters. Finally, the samples were treated with DNase I according to the manufacturer’s instructions. Total RNA was assessed by QuBit® 2.0 Fluorometer (ThermoFisher Scientific) and the associated Qubit RNA assay (Q32852, ThermoFisher Scientific). RNA integrity was confirmed using an Agilent 2100 Bioanalyzer and the associated RNA 6000 pico kit (Agilent Technologies). Samples containing 4.5 μg of total RNA were depleted for ribosomal RNA using a Ribozero rRNA depletion kit (MRZGP126, Cambio), with depletion confirmed by a Bioanalyzer (Agilent). Ion Torrent RNA-Seq library preparation used an Ion total RNA-Seq Kit V2 (4475936, ThermoFisher Scientific) with Ion xpress RNA-Seq BC 01-16 kit barcodes (4475485, ThermoFisher Scientific) as per supplier’s instructions.

### Sequencing and data analysis

The libraries were sequenced using an Ion Torrent PGM (ThermoFisher Scientific) and raw data analysis carried out using the associated Ion Torrent Suite 5.0.2 (ThermoFisher Scientific). Ion PGM template OT2 kits (4480974, ThermoFisher Scientific) were used for emulsion PCR and enrichment steps, with an Ion PGM 200 kit V2 (4482006, ThermoFisher Scientific) for sequencing reactions. Assessment of Ion Sphere Particle quality was undertaken using the Ion Sphere Quality Control Kit (4468656, ThermoFisher Scientific). Triplicate libraries were pooled together to give treated and untreated pools and sequenced initially using Ion 314 chips V2 (4482261, ThermoFisher Scientific). Individual libraries were resequenced using a 318 chip V2 (4484354, ThermoFisher Scientific) to provide technical replicates. All sequence data are deposited on the Sequence Read Archive (SRA) under BioProject: PRJNA603263 (http://www.ncbi.nlm.nih.gov/bioproject/603263). FastQ output files were trimmed (quality score 0.02, discard reads <50 bp in length) and RNA-Seq analysis was carried out using CLC Genomics Workbench version 7.5.1 (Qiagen). The transcriptomics analysis was undertaken using the Empirical Analysis of DGE tool in CLC, with default parameters. The significance of gene expression between the treated and non-treated samples was assessed using Bonferroni adjusted p-value (<0.05) with over two-fold change in expression. Gene comparisons were made using Venn diagrams.

### qRT-PCR

Duplicate samples of total RNA (1 μg) were reverse-transcribed using a qPCRBIO cDNA synthesis kit (PB30.11-02, PCR Biosystems) according to manufacturer’s instructions together with two negative controls (one without reverse transcriptase and another without RNA template). The amount of synthesised cDNA was determined using the QuantiFluor ssDNA system (Promega) with a Qubit® 2.0 Fluorometer (ThermoFisher Scientific) and the samples were stored at −20°C. Primers (Supplementary Table 1) were designed using Primer3 version 0.4.0 software [13, 14] and their concentrations were optimised to decrease primer dimer formation. Amplification reactions were carried out using a 2x qPCRBIO SyGreen Mix Lo-ROX (PB20.11, PCRBiosystems) according to manufacturer’s instructions with 1 μl of cDNA as template. The reactions were run on a Corbett Research 6000 instrument. Amplification efficiency and linearity were determined using a dilution series of DNA. Data analysis was performed on Corbett Rotor Gene 6000 software, Microsoft Office Excel and Minitab statistics package.

### Phenotypic microarray (PM) analysis

Biolog Phenotype MicroArray metabolic panels (PM1, PM2A, PM3B) were used to determine the effect of MGB-BP-3 on growth of *S. aureus* NCTC8325 on single carbon or nitrogen sources. Three dilutions of MGB-BP-3 were tested (0.5 x MIC, 0.25 x MIC, 0.125 x MIC). Biolog chemosensitivity panels (PM11C, PM12B) were used in combination with MGB-BP-3 (0.5 x MIC) to evaluate potential synergy between MGB-BP-3 and other antibiotics. Standard Biolog protocols were used with some variances: sodium pyruvate (5mM, Fisher) was used instead of glucose for PM3B, PM11C and PM12B. Cells were first grown to OD_600_ of 0.3-0.5 and then diluted to OD_600_ 0.03 at inoculation. The various dye mixes were tested and dye mix D was selected. All plates were incubated in the OmniLog instrument (Biolog, Inc. USA) at 37°C for 48 hours. Growth data were recorded every 15 minutes and analysed with OmniLog software Parametric v1.3 and Kinetic v1.3 (Biolog, Inc. USA).

### Melting curve analysis of dsDNA with MGB-BP-3

Three DNA-fragments (70-90 bp long) were amplified by PCR using a standard benchtop PCR machine. The PCR reactions contained *S. aureus* NCTC8325 gDNA as a template, primers (Supplementary Table 1) and GoTaq^®^ G2 Flexi DNA polymerase (M7801, Promega). Resulting PCR products were purified on columns (Isolate II PCR Kit (BIO-52059, Bioline)) that are suitable for >50 bp DNA fragments. The purified DNA fragments were then mixed with 9 μg/ml of MGB-BP-3 in water, and the mixture incubated for 15 min in the dark at room temperature before adding 1:1 volume of ready-made PCR mix that contained an intercalating dye (qPCRBIO SyGreen Mix, PB20.11-05, PCR Biosystems), known not to interfere with PCR. Dissociation-characteristics of dsDNA with MGB-BP-3 were measured and the melting curve analysis was performed by raising temperatures sequentially from 50°C to 99°C using a Rotor-Gene 6000 qPCR machine (RCorbett). Drug-free DNA was used as a control. Statistical significance was assessed by student t-test (n=3).

### Electrophoretic Mobility-Shift assays (EMSA)

The *pdnaD* and *pmraY* promoter fragments were synthesized by Integrated DNA Technologies (Leuven, Belgium) and sub-cloned into plasmid pSR, as a source of DNA fragments for EMSA and footprinting experiments [15]. EMSA was carried out essentially as detailed by Rossiter *et al*. [16]. Purified AatII-HindIII promoter fragments were end-labelled with [γ-^32^P]-ATP and approximately 0.2 nM of each fragment was incubated with varying amounts of MGB-BP-3. The final reaction volume (10 μl) contained HEPES glutamate buffer (pH 8.0, 20 mM HEPES, 5 mM MgCl_2_, 50 mM potassium glutamate, 1 mM DTT, 5% (v/v) glycerol, 0.5 mg/ml BSA) containing 25 μg/ml herring sperm DNA. MGB-BP-3 was incubated with labelled DNA fragments at room temperature for 15 minutes, after which samples were loaded directly onto a running 6% (w/v) polyacrylamide gel (12 v/cm), containing 2% (v/v) glycerol and 0.25 x TBE (Tris/ borate/ EDTA buffer). Gels were analysed using a Bio-Rad Personal Molecular Imager (PMI) and Quantity One software (Bio-Rad).

### DNase I and permanganate footprinting

DNase I and potassium permanganate footprinting experiments were performed on ^32^P-end-labelled AatII-BamHI promoter fragments, using protocols described previously [17, 18]. Each reaction mix (20 μl) contained approximately 1.35 nM template DNA in HEPES glutamate buffer, containing 30 μg/ml herring sperm DNA. DNA fragments were incubated with varying concentrations of MGB-BP-3 for 15 minutes at room temperature before footprinting. For potassium permanganate footprint experiments, herring sperm DNA was omitted from the reaction mixture and either *E. coli* RNA polymerase σ^70^ holoenzyme (Epicentre) or *S. aureus* RNA polymerase σ^A^ holoenzyme [19] were used at a final concentration of 50 nM. The products of all footprinting reactions were analysed by denaturing gel electrophoresis and calibrated with Maxam-Gilbert ‘G+A’ sequencing reactions of the labelled fragment. Gels were quantified using Bio-Rad PMI and Quantity One software.

### Evolution of resistance in the laboratory

A single colony of *S. aureus* (ATCC clone 43300) was transferred to 5 ml L-Broth (LB) and, after overnight incubation at 37° C, 50 μl of the culture was used for serial daily passaging (1/1,000 dilution) in 5 ml LB having either MGB-BP-3 (0.019 μg/ml, 3 replicates), rifampicin (0.4 ng/ml), or no antibiotic (control). After 4 passages (40 generations), concentrations of antibiotics were doubled and passaged daily for 4 further days (40 generations). Aliquots (5 μl) of various 1/10 serial dilutions of the resulting cultures were then spotted in LB-agar plates having either 0.15 μg/ml MGB-BP-3, or 4 ng/ml rifampicin (each corresponding to 2x MIC80), and plates were incubated overnight at 37° C.

## Results

### Exposure to MGB-BP-3 elicits significant changes in the *S. aureus* transcriptome

The minimum inhibitory concentration (MIC) of MGB-BP-3 for *S. aureus* NCTC8325 was 0.19 μg/ml (Supplementary Data MIC). Using fluorescence microscopy, bacterial cells exposed to MGB-BP-3 showed clear fluorescence with no intracellular localisation (Figure 1b). Growth of *S. aureus* in the presence of a sub-lethal concentration of MGB-BP-3 (0.5 x MIC) was found to be affected for 100 minutes post-MGB-BP-3 administration (Figure 1c). Therefore, we decided to perform RNA-Seq experiments at an early time-point (ten minutes) after challenge with MGB-BP-3. This transcriptomic analysis identified 698 transcripts showing significant changes (Supplementary Table 2), some of which were confirmed by quantitative RT-PCR (Supplementary Data qRT-PCR). Previous work by Chaudhuri *et al*. had identified 351 essential genes for growth and survival of *S. aureus* in laboratory culture [20]. Out of the total of 404 downregulated genes in cultures treated with MGB-BP-3, 62 had been classified as essential, and belonged to a variety of gene ontology categories (Figure 2).

**Figure 2:**
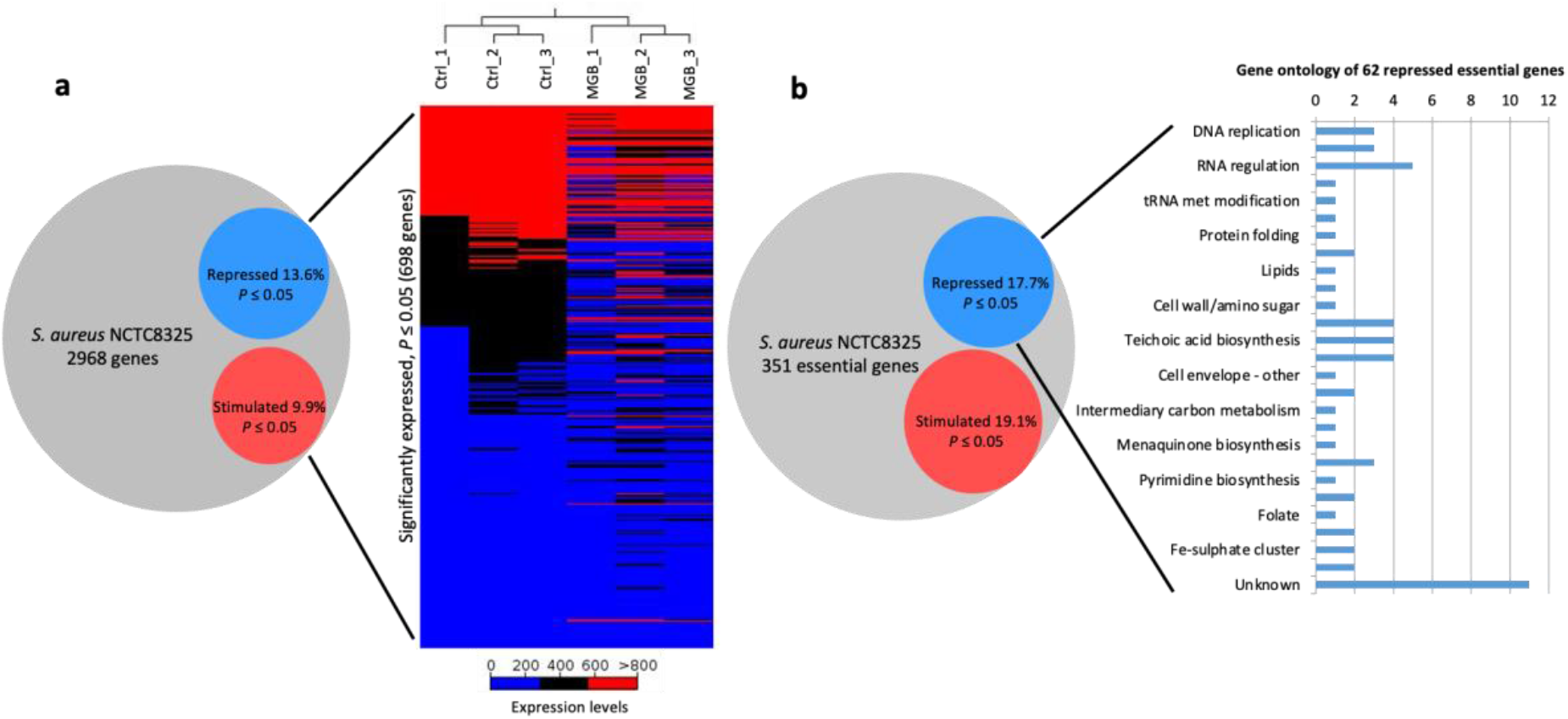
MGB-BP-3 induced transcriptional changes to *S. aureus* NCTC8325 transcriptome on challenge with 0.095 μg/ml MGB-BP-3 (0.5 X MIC) **a**. Venn diagram of RNA-Seq Bonferroni corrected p-values (< 0.05) and a heat map showing hierarchical clustering of the six RNA-Seq samples (3 x ctrl and 3 x drug treated) with a total of 698 significantly expressed genes. **b**. Gene ontology of 62 essential *S. aureus* NCTC8325 genes that were identified in RNA-Seq experiment to be downregulated on challenge with MGB-BP-3.

Further gene ontology analysis of the RNA-Seq data using TIGRFAM protein families [21] identified disruptions in transcription of genes assigned to every Main Role category. Overall, transcription was down-regulated in all categories apart from amino acid biosynthesis, fatty acid and phospholipid metabolism, mobile genetic element functions and transcription (Figures 3 & 4). Transcription of certain non-essential, amino acid biosynthetic genes was stimulated (Supplementary Table 2). Some of these effects were consistent with downregulation of the arginine repressor (SAOUHSC_01617, *argR*) and subsequent consequences on its target genes. Genes involved in biosynthesis of molybdopterin were upregulated, whereas various genes involved in menaquinone and ubiquinone biosynthesis were downregulated significantly (Supplementary Table 2). Some essential genes for pantothenate and coenzyme A biosynthesis and for pyridine nucleotide biosynthesis were downregulated significantly.

**Figure 3:**
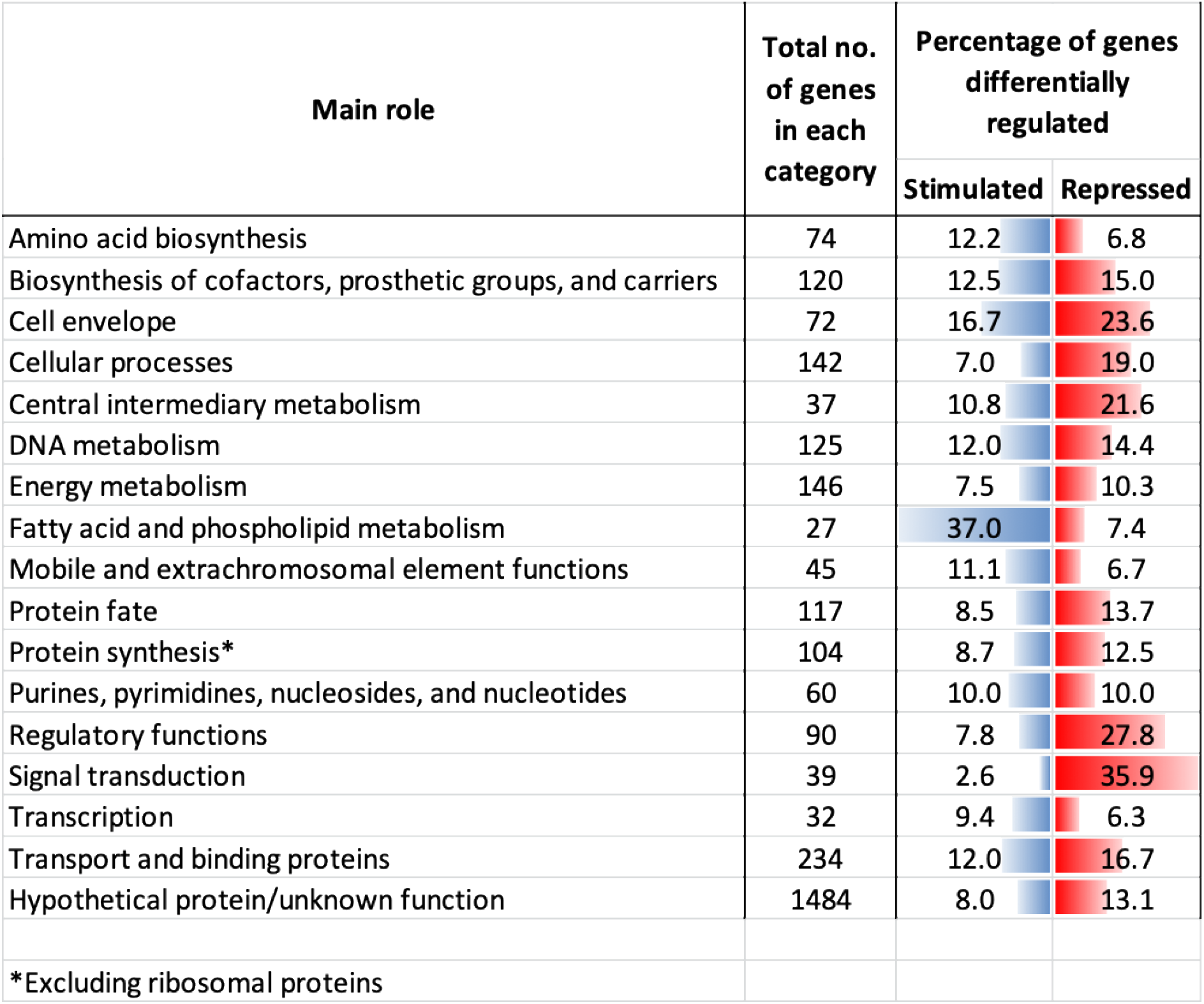
Gene ontology analysis of the *S. aureus* transcriptome after challenge by MGB-BP-3. The percentage of significantly expressed genes belonging to different TIGRFAM protein families is illustrated. The data excludes ribosomal proteins.

**Figure 4:**
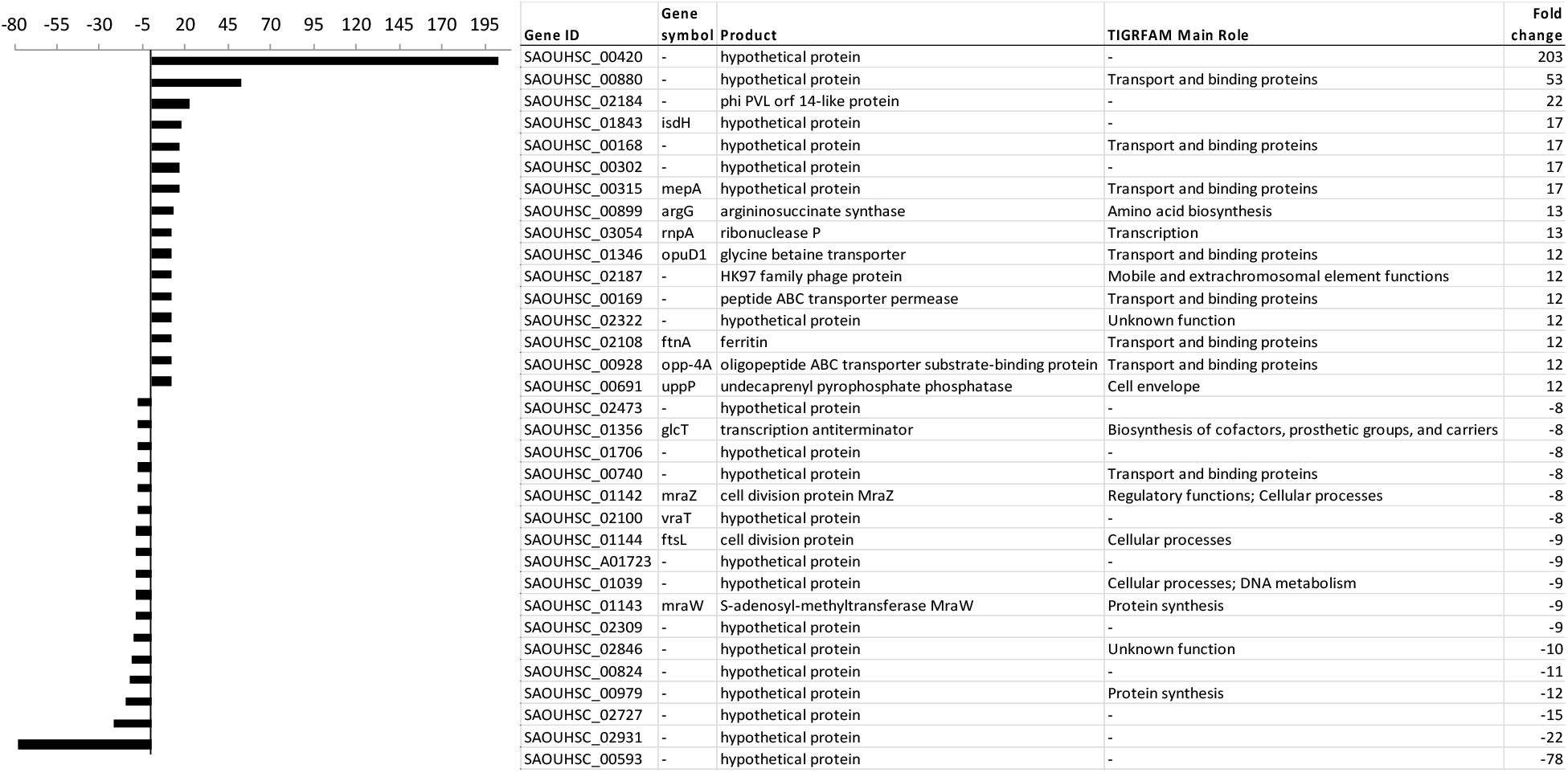
List of genes with the largest fold-changes of the *S. aureus* transcriptome on challenge by MGB-BP-3. The top 10 of genes with the smallest or largest fold-changes are included.

Over half of the genes associated with biosynthesis and degradation of the murein sacculus and peptidoglycan were down-regulated significantly in the RNA-Seq experiment. (Supplementary Table 2). Reduced expression of essential cell division genes) was also noted while the septation ring formation regulator (SAOUHSC_01827, *ezrA*) was also repressed. Some genes involved in DNA replication were down-regulated but, by contrast, some genes involved in DNA supercoiling and primosome formation were up-regulated.

Changes such as those described above could reflect either direct interaction of MGB-BP-3 at the level of the individual gene (considered in the section on *dnaD* and *mraY* promoter regions below), or secondary effects via global regulators. This possibility prompted an assessment of the diverse phenotypes of *S. aureus* when challenged with MGB-BP-3.

### Conditional essentiality allows correlation between phenotype and transcriptotype

Omnilog colorimetric redox phenoarrays were used to directly assess the effect of MGB-BP-3 on metabolism. This approach has the advantage of testing conditional essentiality of down-regulated transcripts in the presence of a sole carbon or nitrogen source. Out of the 191 carbon and 95 nitrogen sources tested and in the absence of MGB-BP-3, *S. aureus* NCTC8325 could utilise 63 carbon and 23 sole nitrogen sources (Supplementary Data; Phenoarrays).

When challenged with sub-MIC levels of MGB-BP-3, *S. aureus* showed several regimes concerning the ability to utilise individual substrates for growth, and in a dose-dependent response (0.5 x MIC, 0.25 x MIC, 0.125 x MIC of MGB-BP-3; Supplementary Data Phenoarrays). Growth on thymidine, glutamine, glycine and arginine as a sole carbon source was affected by MGB-BP-3, whereas growth on ornithine as a single carbon source was not. This was consistent with the results of RNA-Seq (see Discussion).

### Melting curves of MGB-BP-3 bound to dsDNA reveals upstream binding sites

To further identify putative MGB-BP-3 binding sites, upstream regions of two genes shown to be repressed by RNA-Seq (*dnaD* (*SAOUHSC_01470*) and *mraY* (*SAOUHSC_01146*)) were taken forward for melt analysis after being randomly selected from the ten most down-regulated transcripts. MGB-BP-3 would be expected to bind to such a DNA fragment, stabilise its structure, and increase its melting temperature. The genome locations of both genes showed no other genes located directly upstream from the start of the gene of interest. A GC-rich (GC-content 50%) internal region of *gyrA* (*SAOUHSC_00006*) was used as comparative control. The melting temperature of each DNA fragment was increased in the presence of MGB-BP-3, indicating binding of the antibiotic to these AT-rich regions (GC-content of 25-26%) (Figure 5). By contrast, the melting temperature of the control fragment (*gyrA* internal region) was unchanged in the presence of MGB-BP-3.

**Figure 5:**
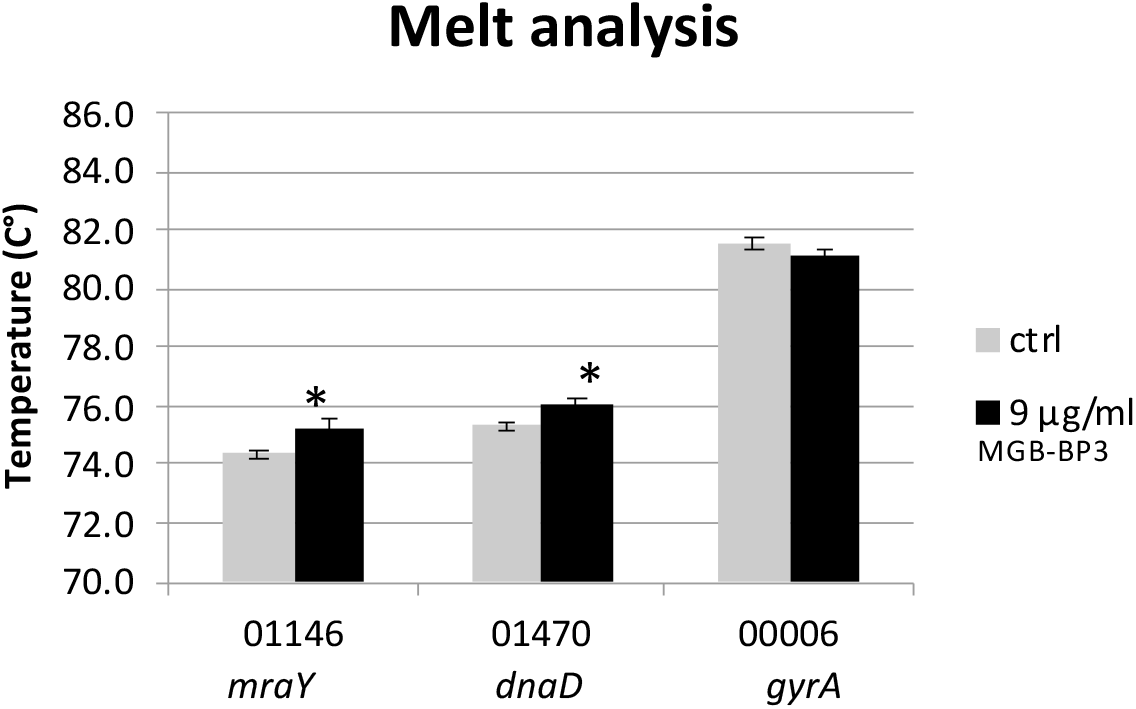
Melt analysis of DNA-MGB-BP-3 (9 μg/ml) complexed with promoter segments of *mraY* and *dnaD*. An AT-rich internal region of *gyrA* was used as DNA segment that should have no binding site for MGB-BP-3. Control samples (grey) did not contain MGB-BP-3. Statistical significance (p ≤ 0.05) is indicated with an asterisk. Error bars, s.d., *n* = 3

### MGB-BP-3 binds to the *dnaD* and *mraY* promoter regions to interfere with transcriptional initiation

DNA melt curves indicated that MGB-BP-3 binds to DNA with some specificity. To examine this further, EMSA assays investigated the binding of MGB-BP-3 to end-labelled DNA fragments, which carried the *dnaD* and *rmaY* promoter regions (*pdnaD* and *pmraY*) (Figure 6). These were chosen as representative of genes that were shown to be down-regulated on challenge with MGB-BP-3 by RNA-Seq and qPCR.

**Figure 6:**
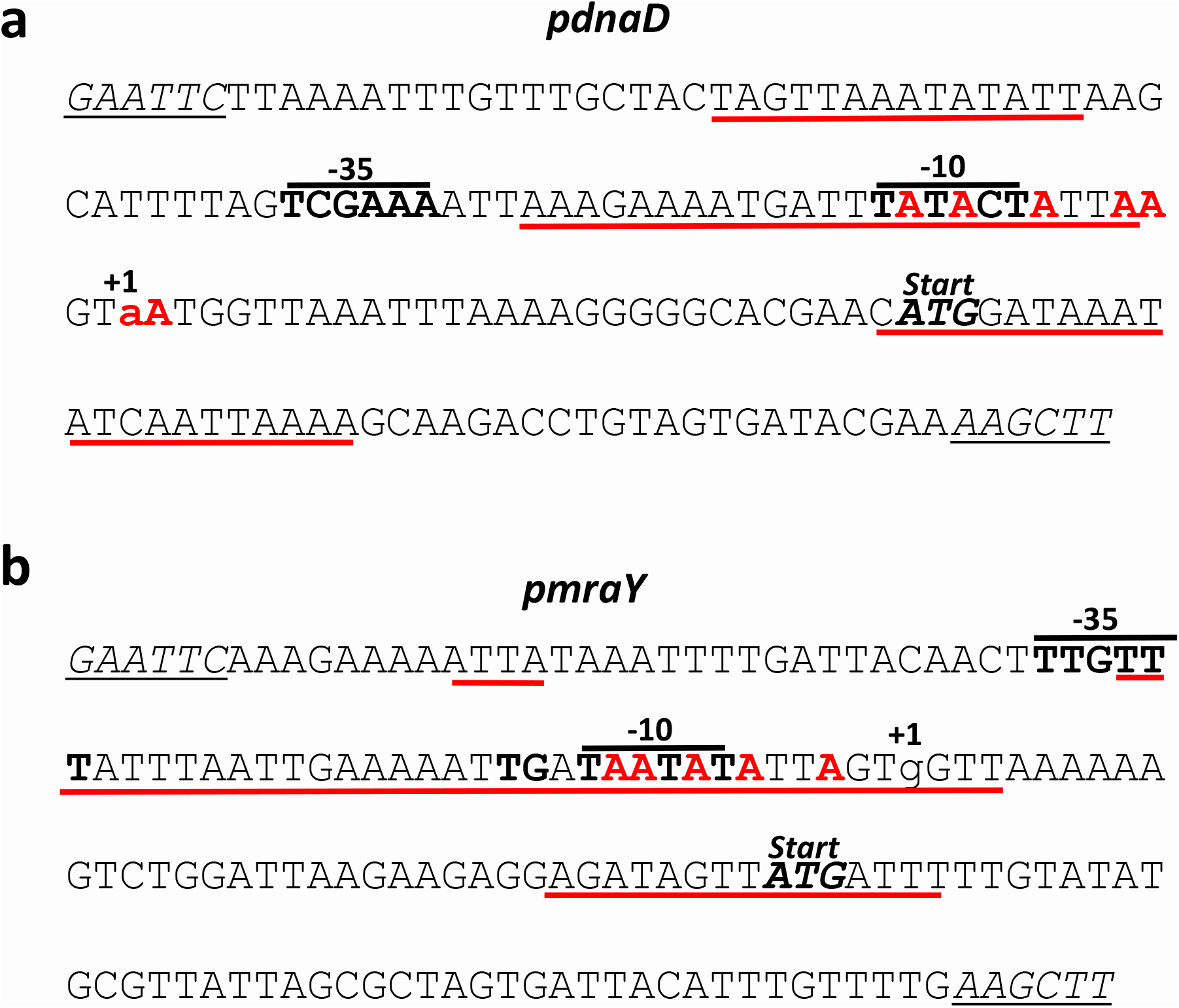
Organisation of the *pdnaD* (a) and *prmaY* (b) promoter regions of *S. aureus*. (+1 and lower case) denotes the experimentally determined transcription start site [27, 28] and the predicted −10 and −35 regions are also shown in bold [29]. The cleavage sites produced by potassium permanganate footprinting are shown in red and bold, whilst the extent of the high affinity MGB-BP-3 binding sites, determined by DNase I footprinting, are underlined in red. The translation start of each gene (ATG) is italicised and bold. The EcoRI and HindIII sites introduced onto each fragment to facilitate cloning into pSR are shown italicised and underlined.

MGB-BP-3 is a small molecule (MW 746 Da). A ladder of fragments with altered mobility was observed for each, indicating that each promoter fragment carried multiple binding sites for MGB-BP-3 (Figure 7). Thus, to pinpoint these MGB-BP-3 binding sites at high resolution, DNase I footprinting was used (Figure 8). For each promoter fragment, discrete high affinity binding sites were observed at low concentrations of MGB-BP-3, whilst additional protections from DNase I were observed as the concentration of antibiotic was increased. In particular, high affinity binding sites for MGB-BP-3 correlated with regions of A/T richness, close to the experimentally determined transcription start sites and overlapping the predicted −10 elements of each promoter (Figures 6 and 8) [27–29]. As positioning of such high affinity binding sites for MGB-BP-3 could interfere with transcriptional initiation, potassium permanganate footprinting was used to examine promoter isomerisation by RNA polymerase at each promoter. Single-stranded DNA, generated by DNA melting during transcriptional initiation, is sensitive to modification by permanganate, which can be detected by gel electrophoresis [18, 30]. In the absence of MGB-BP-3, both *E. coli* and *S. aureus* RNA polymerase holoenzyme could recognise *pdnaD* and *pmraY*, unwinding the DNA surrounding the transcription start site and each −10 promoter element. However, in the presence of MGB-BP-3, unwinding around the promoter was completely inhibited (Figures 9 and 10), confirming that MGB-BP-3 prevents transcription initiation at *pdnaD* and *pmraY* by occluding the promoter region and preventing RNA polymerase isomerisation.

**Figure 7:**
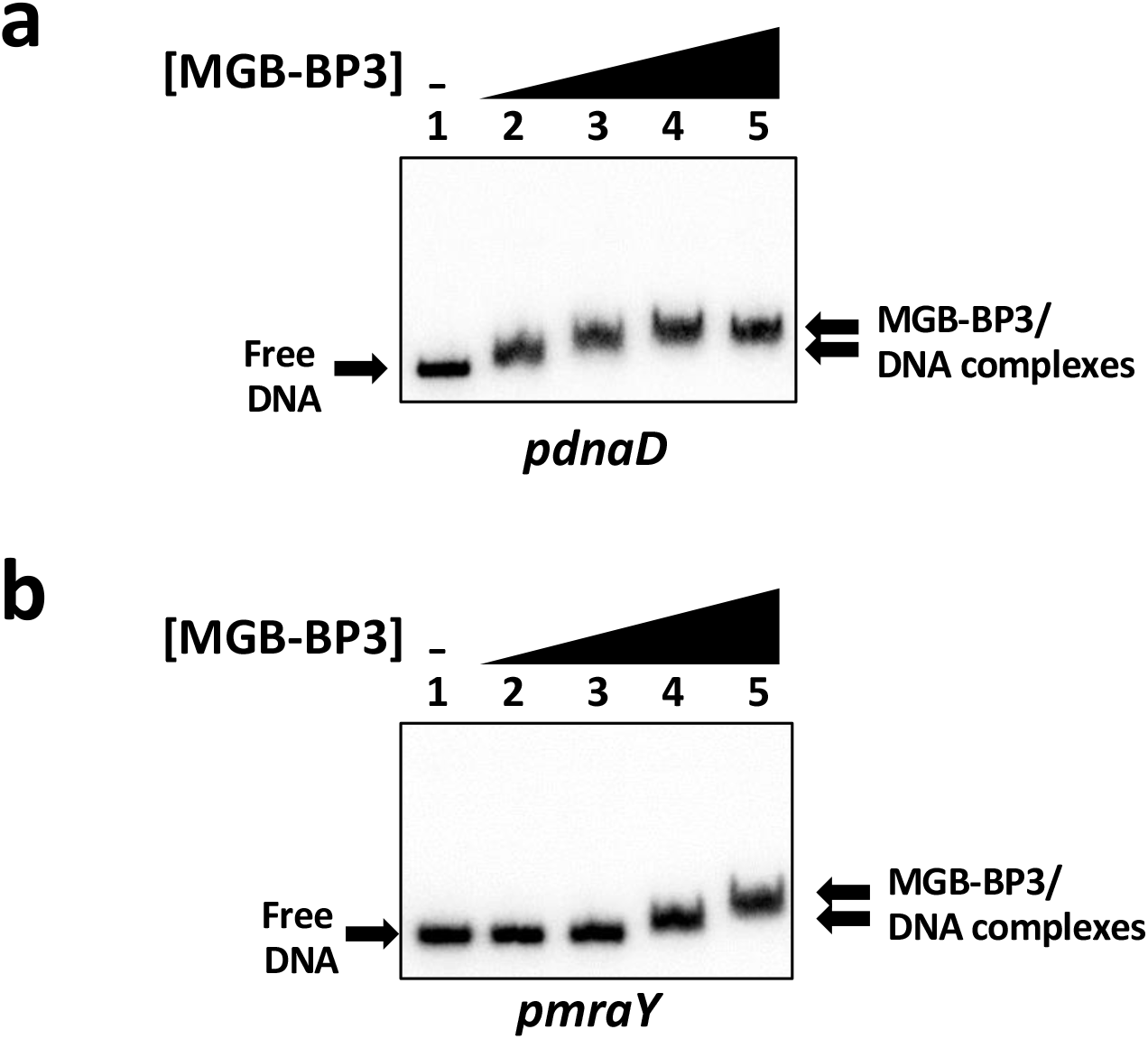
MGB-BP-3 binds to the *pdnaD* and *prmaY S. aureus* promoters. EMSA of the (a) *pdnaD* and (b) *pmraY* promoter regions with MGB-BP-3. End-labelled AatII-HindIII fragments were incubated with increasing concentrations of MGB-BP-3 as follows: lane 1, no MGB-BP-3; lane 2, 1.25 μg/ml; lane 3, 2.5 μg/ml; lane 4, 5 μg/ml; lane 5, 10 μg/ml.

**Figure 8:**
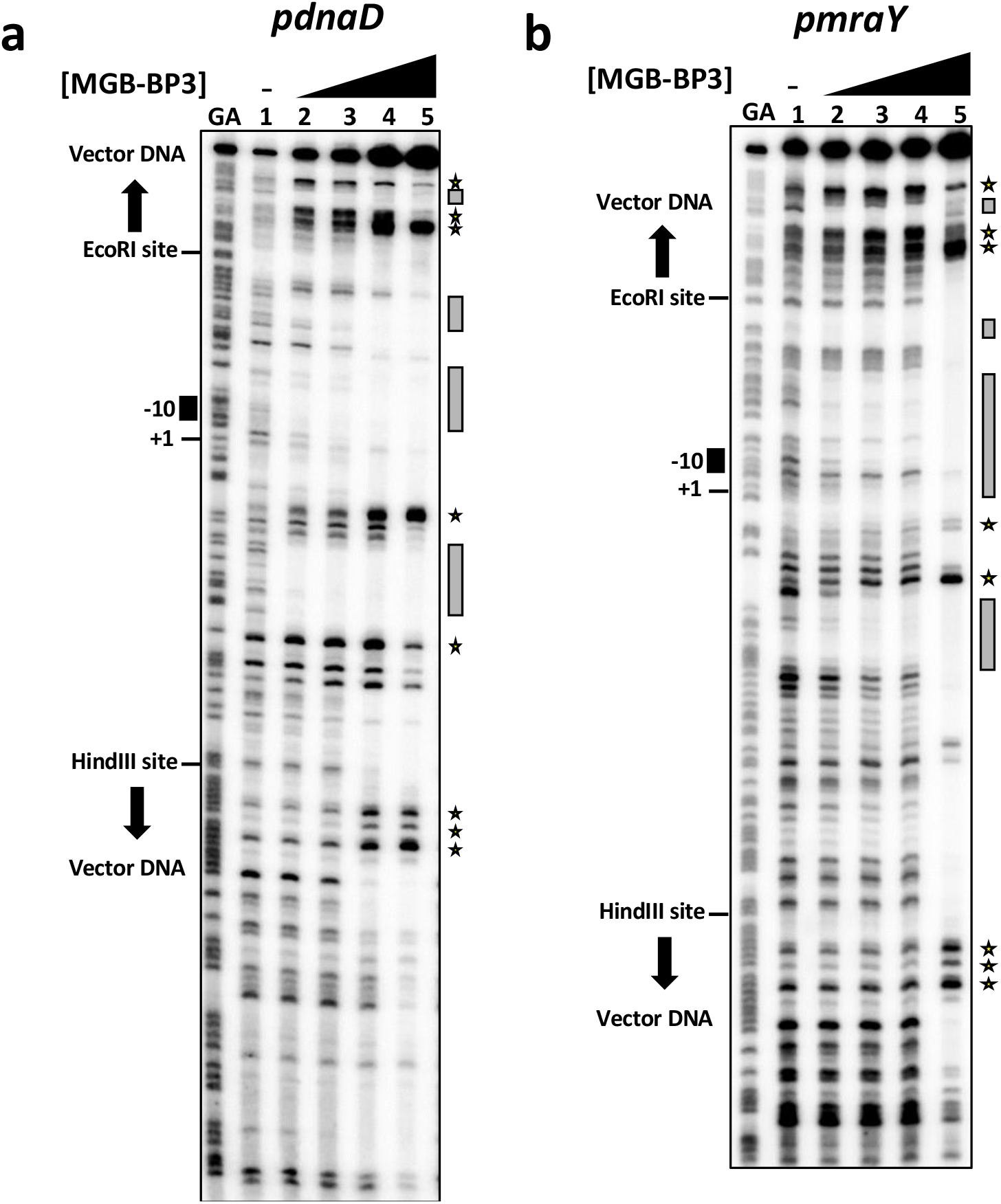
MGB-BP-3 binds to discrete sites within the *pdnaD* and *prmaY S. aureus* promoters. DNase I footprinting analysis of the (a) *pdnaD* and (b) *pmraY* promoter regions when bound by MGB-BP-3. End-labelled AatII-BamHI fragments were incubated with increasing concentrations of MGB-BP-3 and subjected to DNase I footprinting. The concentrations of MGB-BP-3 used were as follows: lane 1, no MGB-BP-3; lane 2, 1.25 μg/ml; lane 3, 2.5 μg/ml; lane 4, 5 μg/ml; lane 5, 10 μg/ml. Gels were calibrated using Maxam-Gilbert ‘G+A’ sequencing reactions (lane GA) and the location of the −10 element and transcription start site (+1) for each promoter is indicated. Regions of high affinity MGB-BP-3 protection are indicated by grey boxes and hypersensitive sites produced by MGB-BP-3 binding are labelled with stars. The locations of the EcoRI and HindIII sites and the pSR vector sequences on each AatII-BamHI fragment are also marked.

**Figure 9:**
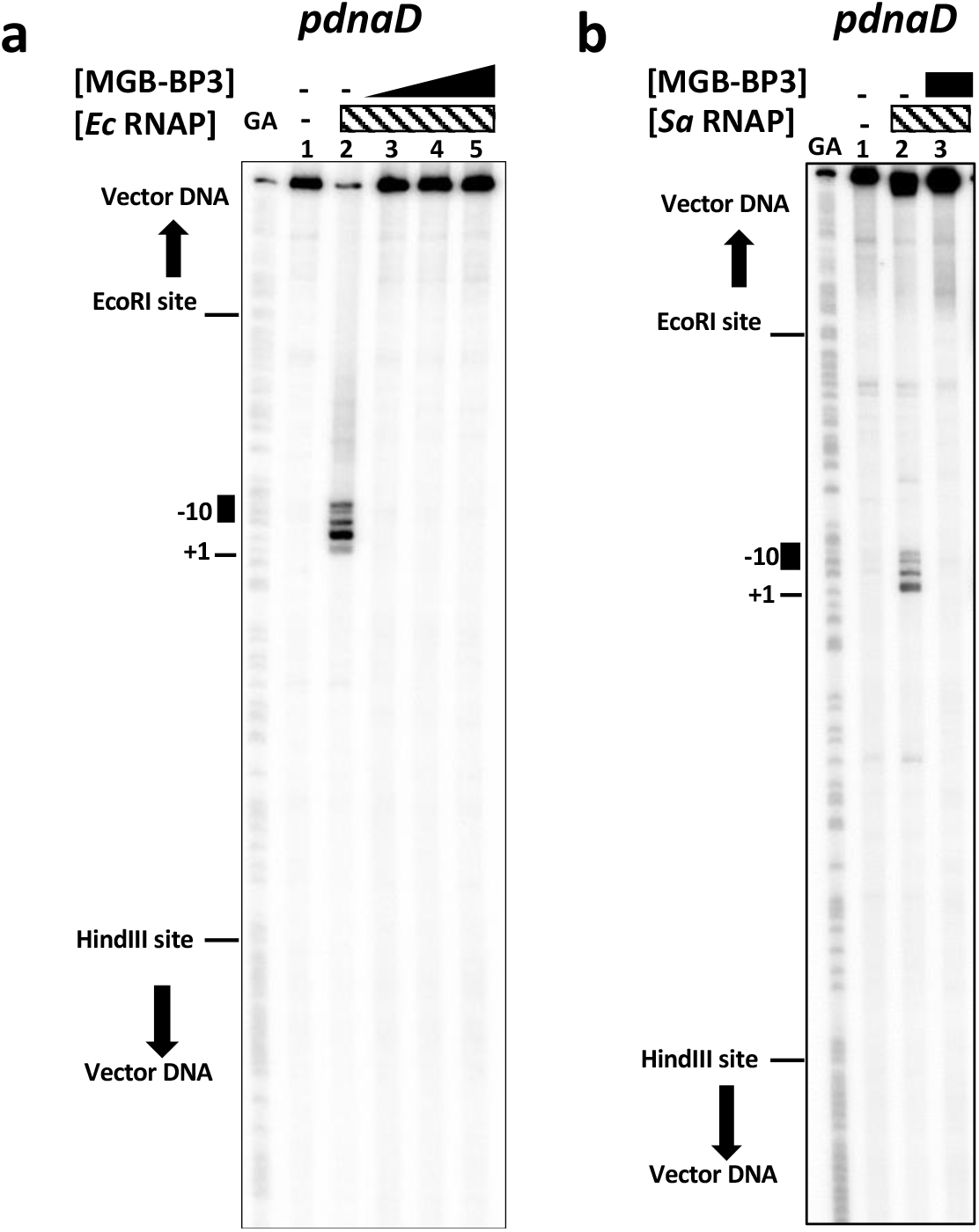
MGB-BP-3 prevents promoter unwinding by both *E. coli* and *S. aureus* RNA polymerase holoenzyme at the *pdnaD S. aureus* promoter. Potassium permanganate footprinting analysis of the *pdnaD* promoter region when bound by MGB-BP-3. (a) End-labelled AatII-BamHI *pdnaD* promoter fragment was pre-incubated with MGB-BP-3, challenged with 50 nM *E. coli* RNA polymerase holoenzyme and subjected to potassium permanganate footprinting. The concentrations of MGB-BP-3 were as follows: lanes 1 and 2, no MGB-BP-3; lane 3, 1.25 μg/ml; lane 4, 2.5 μg/ml; lane 5, 5 μg/ml. (b) End-labelled AatII-BamHI *pdnaD* promoter fragment was pre-incubated with MGB-BP-3, challenged with 50 nM *S. aureus* RNA polymerase holoenzyme, saturated with σ^A^, and subjected to potassium permanganate footprinting. The concentrations of MGB-BP-3 were as follows: lanes 1 and 2, no MGB-BP-3; lane 3, 2.5 μg/ml. Gels were calibrated using Maxam-Gilbert ‘G+A’ sequencing reactions (lane GA) and the location of the predicted *pdnaD* −10 element and experimentally determined transcription start site (+1) is indicated [28]. The positions of the EcoRI and HindIII sites and the pSR vector sequences on each AatII-BamHI fragment are also marked.

**Figure 10:**
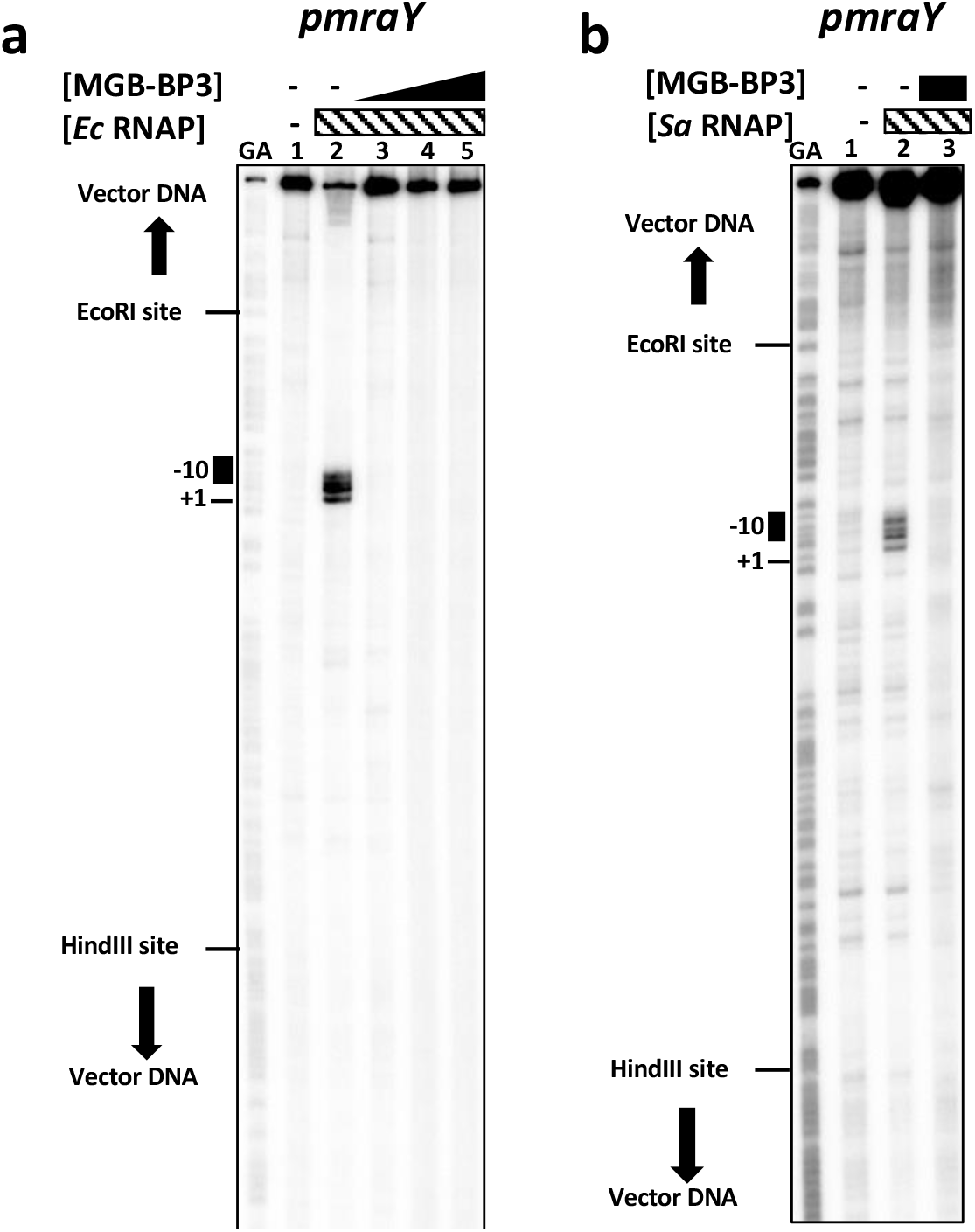
MGB-BP-3 prevents promoter unwinding by both *E. coli* and *S. aureus* RNA polymerase holoenzyme at the *pmraY S. aureus* promoter region. Potassium permanganate footprinting analysis of the *pmraY* promoter region when bound by MGB-BP-3. (a) End-labelled AatII-BamHI *pmraY* promoter fragment was pre-incubated with MGB-BP-3, challenged with 50 nM *E. coli* RNA polymerase holoenzyme and subjected to potassium permanganate footprinting. The concentrations of MGB-BP-3 were as follows: lanes 1 and 2, no MGB-BP-3; lane 3, 1.25 μg/ml; lane 4, 2.5 μg/ml; lane 5, 5 μg/ml. (b) End-labelled AatII-BamHI *pmraY* promoter fragment was pre-incubated with MGB-BP-3, challenged with 50 nM *S. aureus* RNA polymerase holoenzyme, saturated with σ^A^, and subjected to permanganate footprinting. The concentrations of MGB-BP-3 were as follows: lanes 1 and 2, no MGB-BP-3; lane 3, 2.5 μg/ml. Gels were calibrated using Maxam-Gilbert ‘G+A’ sequencing reactions (lane GA) and the predicted location of the *pmraY* −10 element and the experimentally determined transcription start site (+1) is indicated [27]. The positions of the EcoRI and HindIII sites and the pSR vector sequences on each AatII-BamHI fragment are also marked.

### Can *S. aureus* evolve to be resistant to MGB-BP-3?

As MGB-BP-3 binds to multiple sites in the bacterial genome, with consequential reduction in transcription of essential genes, a great number of mutations would have to occur within a short time period to result in resistance to this antibiotic. This contrasts with other antibiotics such as rifampicin, which bind to a single target, such that a single genetic mutation is sufficient to confer resistance. Indeed, after serial passaging of three independent populations of *S. aureus* at sub-MIC80 concentrations of MGB-BP-3 (up to 0.5x MIC80) for 80 generations, no resistant clone was isolated. By contrast, serial passaging for the same period at sub-MIC80 concentrations of rifampicin (up to 0.4x MIC80), resistant colonies were identified even at the lowest spotted concentration (corresponding to OD_600_ 10^−8^, Supplementary Data; Resistance). In a second experiment, a population of *S. aureus* was passaged at 0.66x MIC80 of MGB-BP-3 for 280 generations, but no resistance was observed (data not shown).

## Discussion

DNA minor groove-binding drugs have shown promise in the treatment of a variety of diseases [31]. Amongst these, novel antibiotic MGB-BP-3 is in development for treatment of *Clostridioides difficile* infections and has completed Phase II clinical trials successfully. Here, *S. aureus* was used as a model organism to study MGB-BP-3 mode of action, since much of the preliminary work for the MGB-BP-3 development program had already been done with this organism. Moreover, this choice simplified the growth conditions for the Biolog phenotypic array work.

Transcriptomics identified 62 essential genes that were repressed in the presence of this DNA-binding antibiotic (Figure 2). Transcriptional changes in seven of these genes (*argH, cdd, citC, gapA, hup, mvaK2* and *pyrF*) was confirmed independently by qRT-PCR. Several genes with regulatory functions that could account for some of the changes seen in the RNA-Seq data had expression altered significantly on exposure to MGB-BP-3. (Supplementary Table 2). For instance, anti-anti-sigma factor *rsbV* (SAOUHSC_02300) was repressed whereas RNA polymerase sigma factor RpoD coded by *sigA* (SAOUHSC_01662) was stimulated. Moreover, two-component systems such as *walKR* (SAOUHSC_00021 and SAOUHSC_00020) and *phoPR* (SAOUHSC_01800 and SAOUHSC_01799), together with glutamine synthetase repressor *glnR* (SAOUHSC_01285), DNA-binding response regulator *srrA* (SAOUHSC_01586), transcriptional regulator *nrdR* (SAOUHSC_01793) and transcription of LexA repressor (SAOUHSC_01333, *lexA*) were down-regulated significantly.

Microtiter plate-based phenoarrays offer an alternative strategy to investigate challenge of bacterial cells by MGB-BP-3, illustrating the conditional essentiality effects of the antibiotic and providing data (Supplementary Data Phenoarrays) to compare with the transcriptomic response. Thus, none of the genes for ornithine metabolism (SAOUHSC_00076, SAOUHSC_00148, SAOUHSC_00150, SAOUHSC_00894, SAOUHSC_01128, SAOUHSC_02967, SAOUHSC_02968) had significantly altered expression profiles on challenge by MGB-BP-3, consistent with the data from the phenotypic arrays. Although arginine is a precursor of ornithine [25], its metabolism was affected negatively on challenge by MGB-BP-3 in a dose dependent manner. Transcriptomics revealed that expression of *ahrC* (SAOUHSC_01617) was significantly downregulated (by 6-fold) on exposure to MGB-BP-3. AhrC acts as a repressor of arginine biosynthesis and an activator of the arginine catabolic pathway in *B. subtilis* [26]. Thus down-regulation of arginine catabolism directly could explain the differences seen in response to MGB-BP-3 for growth on either arginine or ornithine as a single carbon source. Further correlations between the transcriptomic data and phenotypic arrays reinforced the multiplicity of effects on the bacterial cells on challenge with MGB-BP-3. Taken together, our data supports the hypothesis that MGB-BP-3 binds to and inhibits multiple essential promoters on the *S. aureus* chromosome.

Reflecting the current annotation status of *S. aureus,* genes encoding hypothetical proteins and proteins with unknown function were the largest category in the transcriptomic data that showed a general trend of transcriptional repression. By contrast, hypothetical protein SAOUHSC_00420 was the most upregulated gene (203-fold) in the RNA-Seq dataset (Figure 4). It is predicted to encode a sodium-dependent transporter that has been implicated in extracellular DNA release during biofilm formation in *S. aureus* [24]. The second most enhanced gene *SAOUHSC_00880* (53-fold change) was a hypothetical protein, also assigned to the category of transport and binding proteins. A further 7 out of 16 of the most upregulated genes belonged to this category, which may reflect a generalised antibiotic stress response.

Direct inhibition of transcription at individual gene loci by was investigated further by focussing on two essential genes as exemplars, *dnaD* (SAOUHSC_01470) and *mraY* (SAOUHSC_01146). DNA melt curves demonstrated binding of MGB-BP-3 to upstream regions of these genes (Figure 5), which was confirmed by EMSA (Figure 7). Much higher resolution of MGB-BP-3 binding upstream of these genes was determined with DNAse I footprinting assays (Figures 6 and 8) demonstrating that high affinity binding sites for MGB-BP-3 overlap both the *dnaD* and *mraY* promoters. The presence of multiple MGB-BP-3 binding sites upstream of both *dnaD* and *mraY* may explain the strength of the inhibitory effect at these particular promoters (Figures 6 and 8). Permanganate footprinting of these promoters in the presence of purified RNA polymerase (RNAP) from both *E. coli* and *S. aureus* demonstrated that MGB-BP-3 binding to the −10 element inhibited the isomerisation of the promoter by RNAP holoenzyme (core polymerase plus SigA, Figures 6, 9 and 10). Specificity of transcription from promoters such as *dnaD* and *mraY* is conferred by the SigA subunit of RNAP. Thus, it has been demonstrated that MGB-BP-3 binds to certain SigA-dependent promoter regions, preventing transcription of these genes. This is almost certainly not the sole mechanism of action of MGB-BP-3 since transcription is not the only essential protein-DNA interaction, but is sufficient across the various target genes to explain the catastrophic death on the bacterial cells on challenge with MGB-BP-3.

Microbial drug resistance has become a global health problem and the need to bring new antibiotics to the market is increasing [32]. Resistance against new drugs can develop within the first couple of years of the drug entering the market [33]. Our data suggests that MGB-BP-3 binds to multiple sites on *S. aureus* chromosome, making it less likely for resistance to evolve by drug target mutation. To test this hypothesis, laboratory-directed evolution experiments using sub-inhibitory concentrations of MGB-BP-3 showed no resistance in *S. aureus*. In contrast, resistance to the single target RNA-polymerase inhibitor rifampicin arose rapidly (Supplementary Data; Resistance).

In summary, a novel mode of action for MGB-BP-3 against *S. aureus* has been demonstrated by thorough investigation of two exemplar drug-binding sites on the *S. aureus* chromosome. RNA-Seq analysis revealed a total of 698 transcripts with drug altered expression profiles. It is, therefore, highly likely that there are further multiple MGB-BP-3 binding sites on the *S. aureus* chromosome. Further analysis of DNA binding site sequences combined with rational design of other antibiotic MGBs could provide a powerful methodology for new drug development.

## Supporting information

Supplementary data; Minimal Inhibitory Concentrations

Supplementary data; Phenoarrays

Supplementary data; qRT-PCR

Supplementary data; Resistance

## Acknowledgements

LK was supported by BBSRC grant BB/K019600/1 to NPT, ISH and CJS. Work in NPT’s laboratory is also supported by BBSRC grants BB/V509243/1 and BB/S507106/1. KL was supported by CSO grant TCS/16/24 to ISH, CJS and NPT. DFB was supported by BBSRC grant BB/R017689/1. We thank Ramesh Wigneshweraraj and Lynn Burchell for kindly providing purified *S. aureus* core RNA polymerase and σ^A^ protein.

## Author Contributions

NPT conceived all experiments, LK conceived and performed all microscopy, RNA-sequencing, qRT-PCR, melting analysis and phenotypic microarray experiments, DFB conceived and performed all the EMSA, DNase I and permanganate footprinting experiments, KL performed evolution of resistance experiment and TS performed independent growth studies using single carbon source, LK and NPT analysed data and wrote the draft manuscript. DFB, KL, ISH and CJS analysed data and contributed to the manuscript. NPT, ISH and CJS acquired funding.

## Supplementary information

**Supplementary Data –MIC. Comparison of the MIC of vancomycin and MGB-BP-3 against***S. aureus*

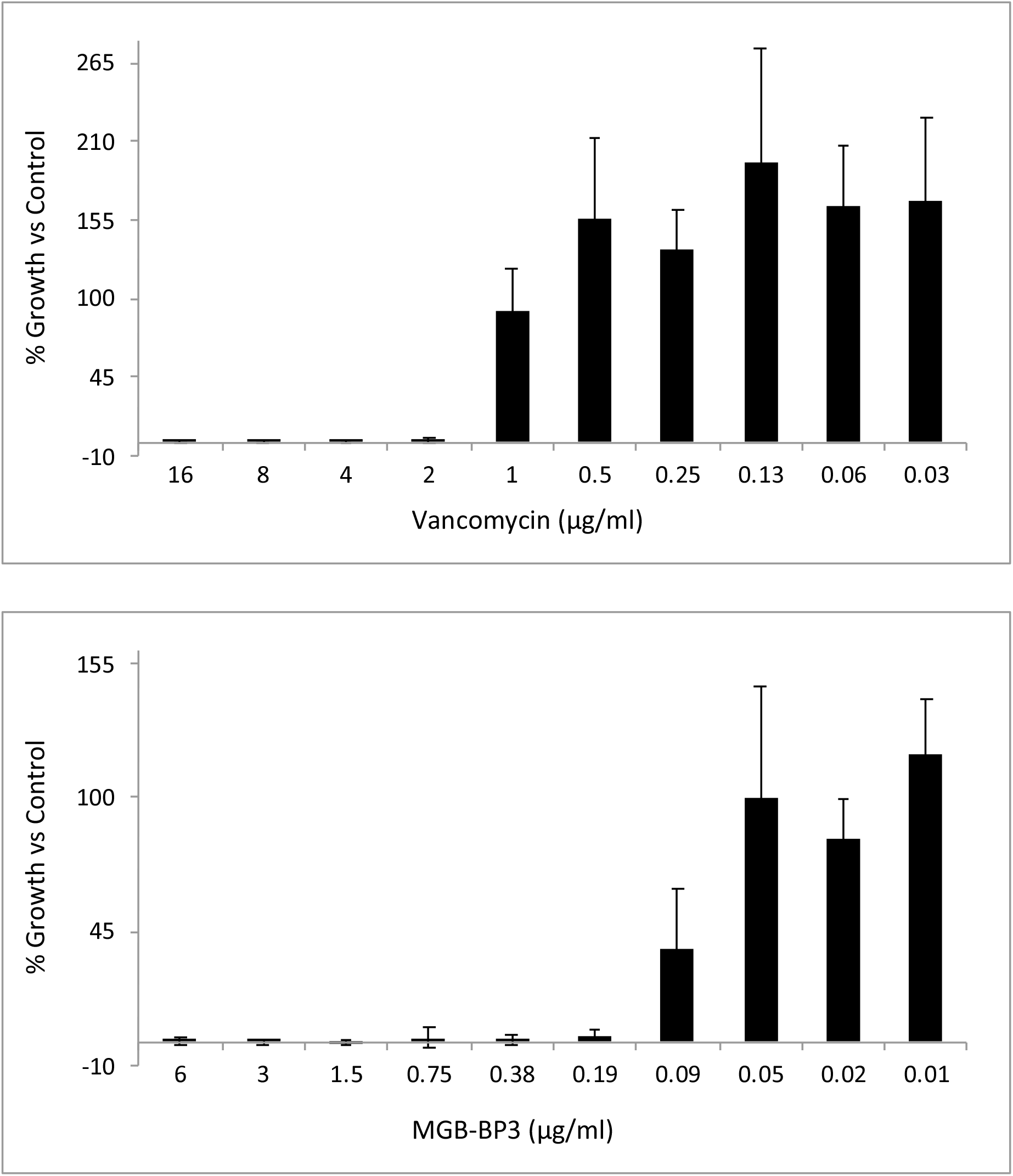

**Supplementary Data – qRT-PCR. Confirmation of selected transcriptional effects of MGB-BP-3 by qRT-PCR.**

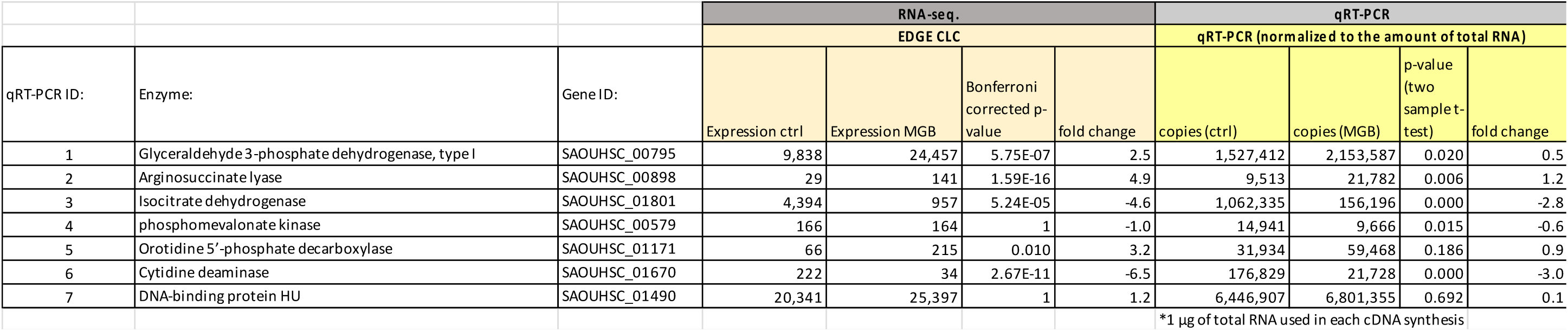

**Supplementary Data – Phenoarrays.**

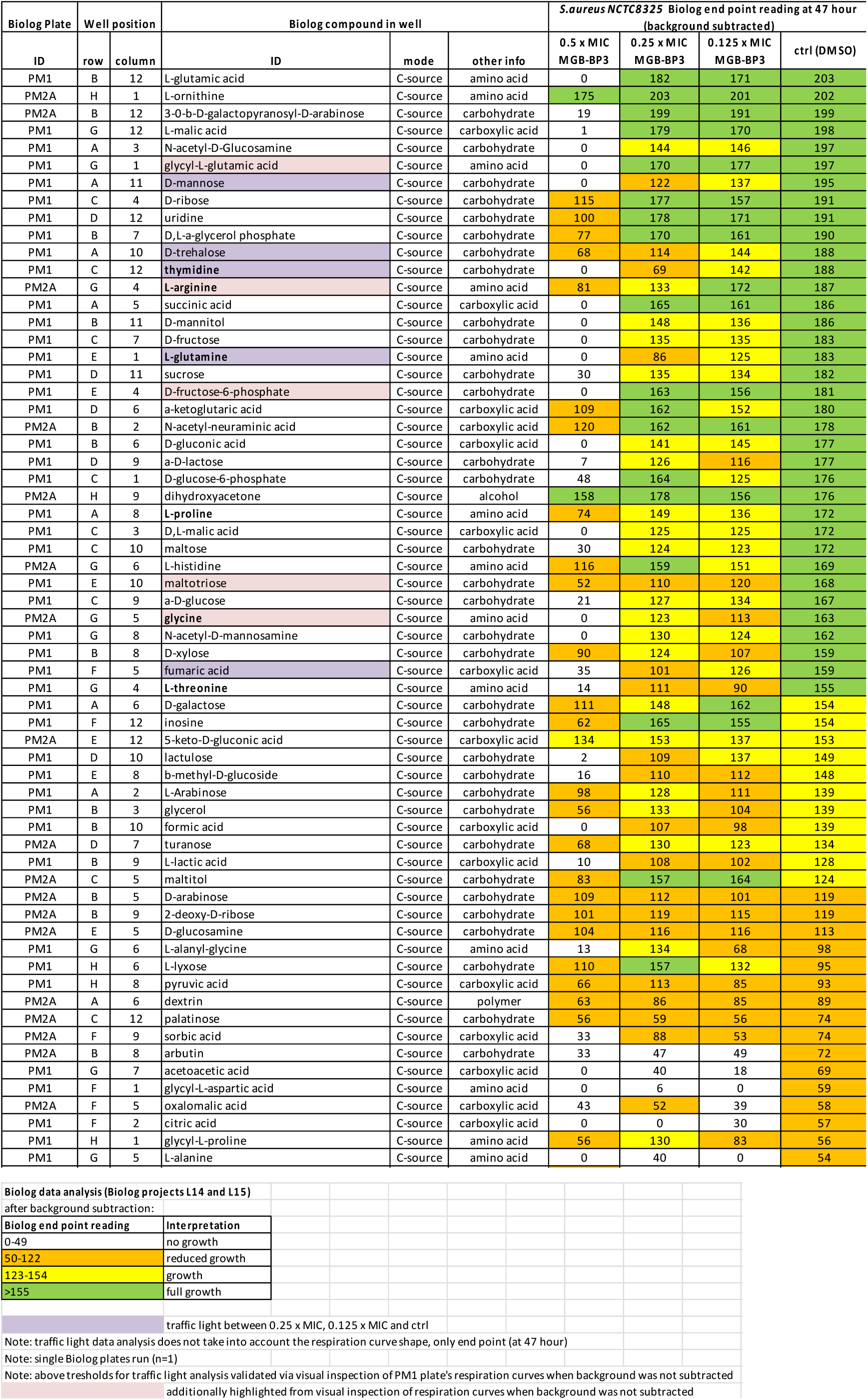

**Supplementary Data – Resistance. Assessment of S. aureus resistance to MGB-BP-3 following serial passaging in LB containing either MGB-BP-3 (3 independent populations at up to 0.5x MIC80), rifampicin (one population at up to 0.4x MIC80), or no antibiotic (control) for 80 generations. Cultures at corresponding OD600 were spotted on agar plates containing either rifampicin or MGB-BP-3 at 2x MIC80 concentrations.**

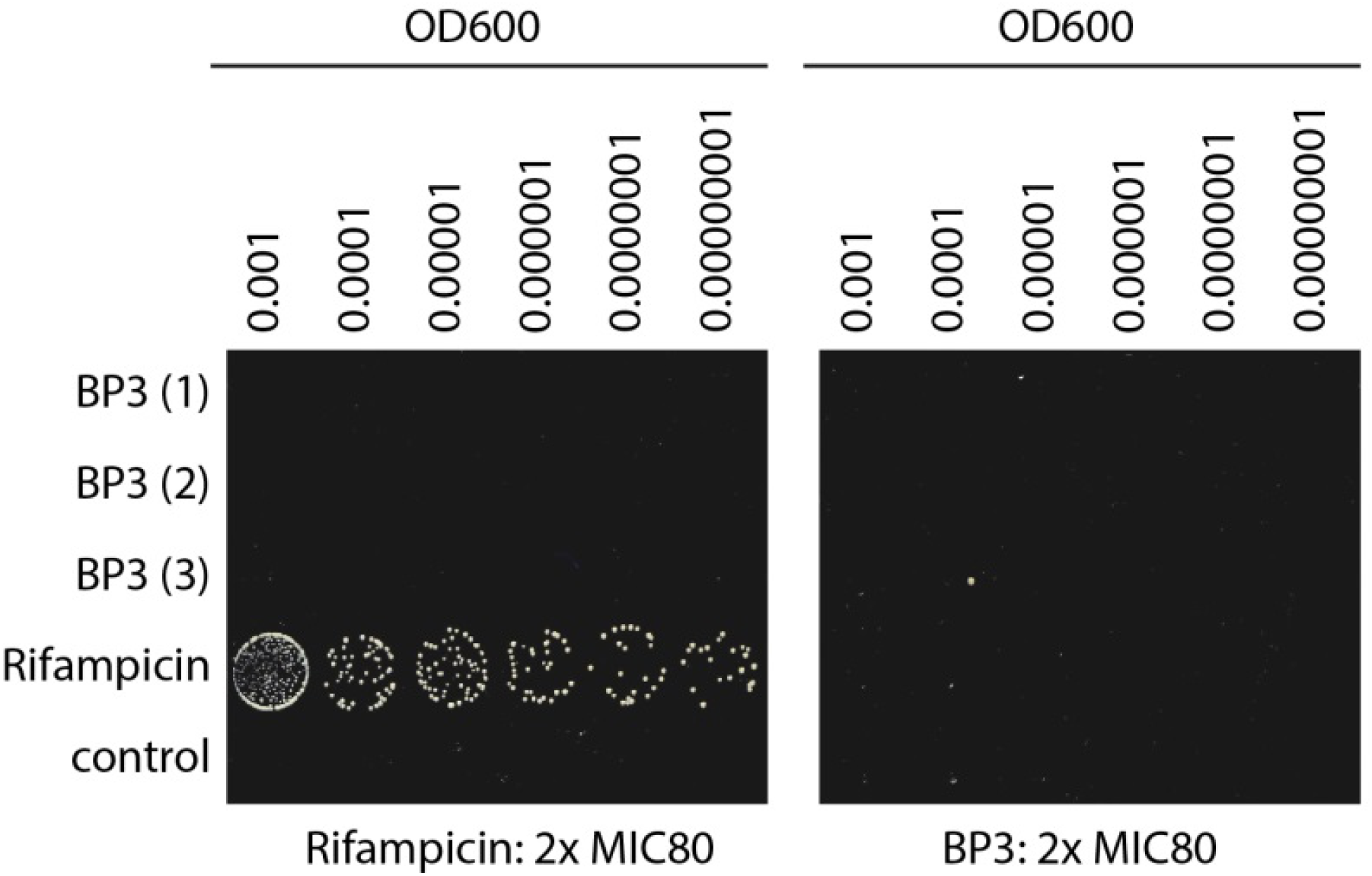

---

**Supplementary Table 1:**
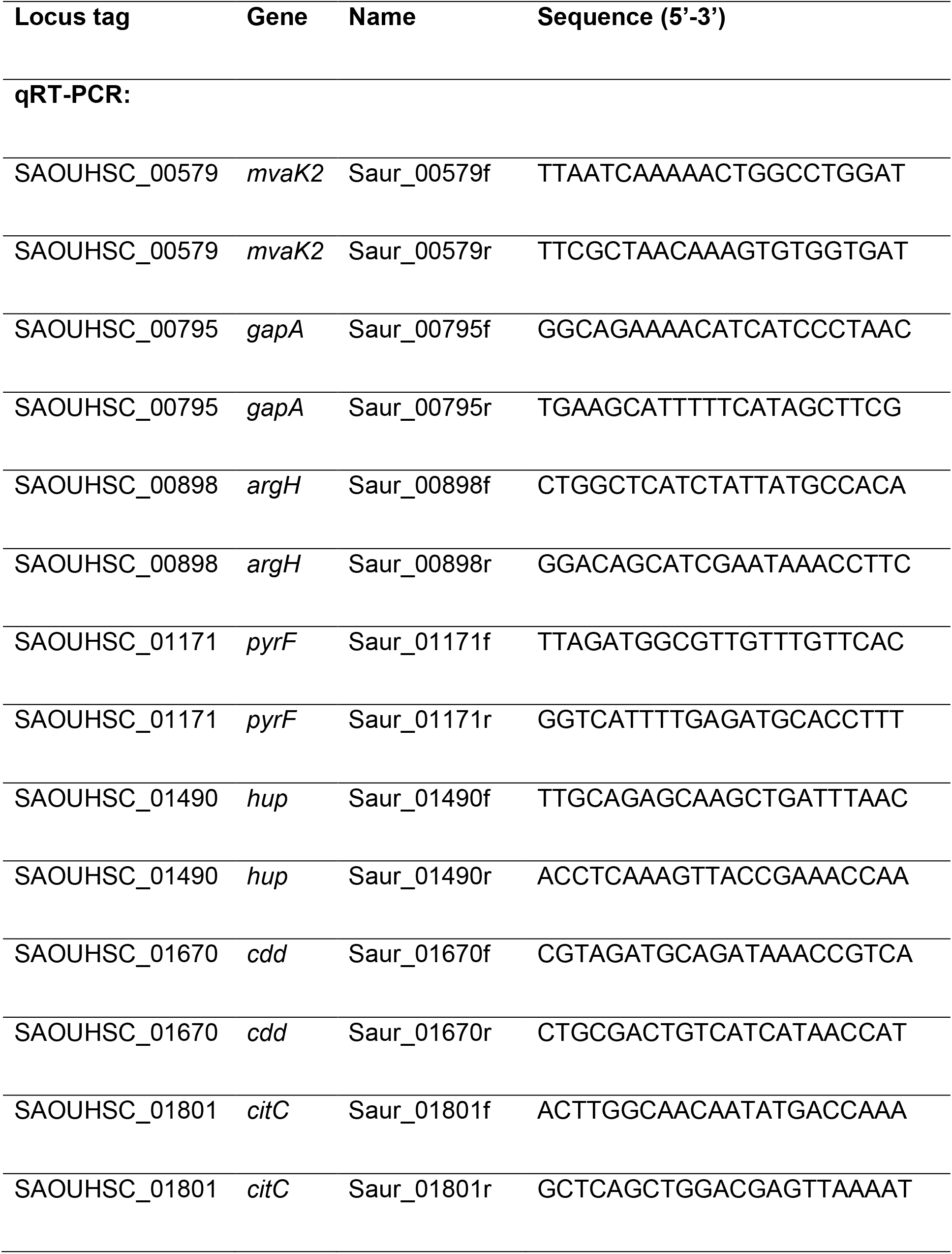
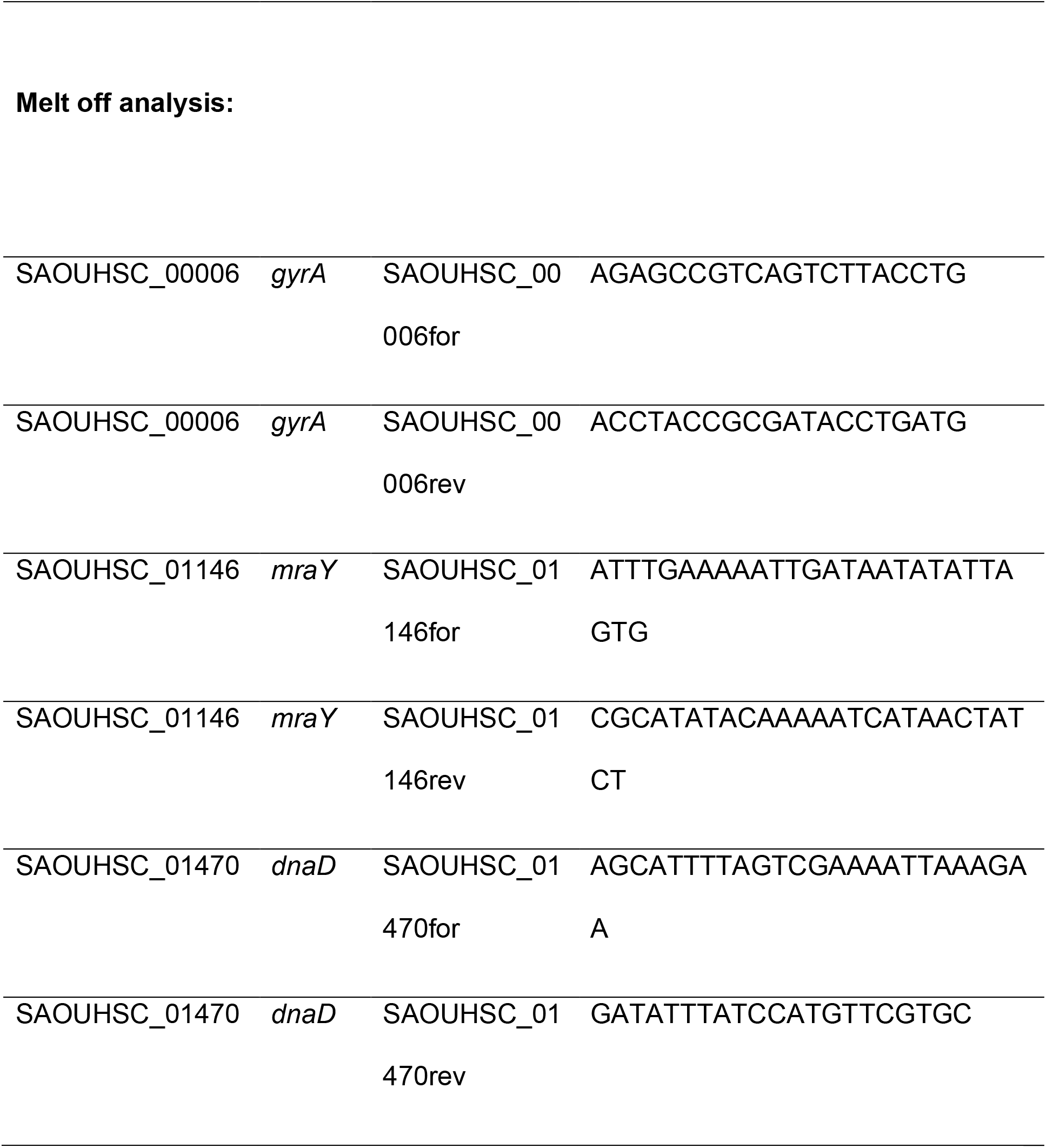
Primers used in qRT-PCR expression studies and Melt off analysis

**Supplementary Table 2.**
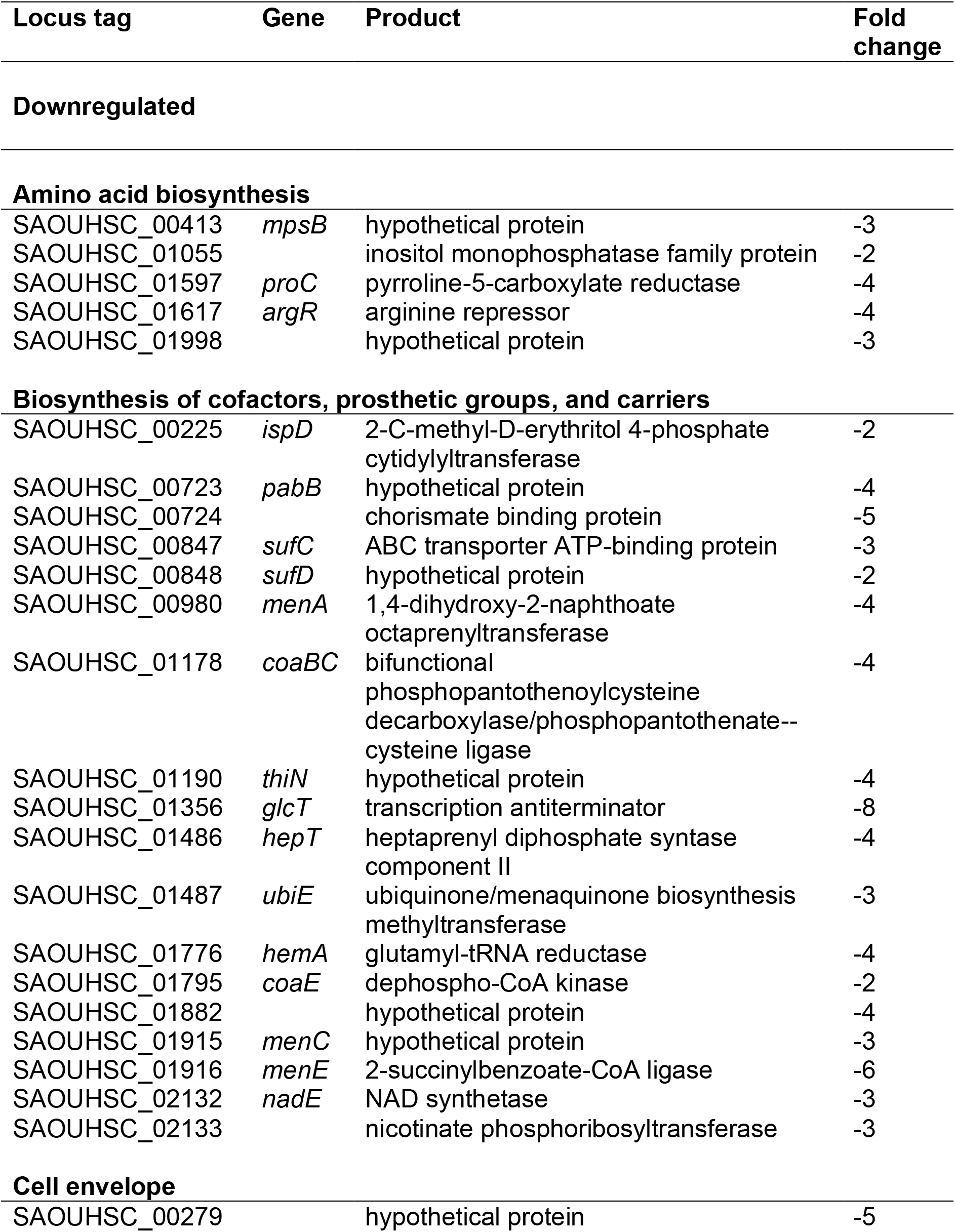
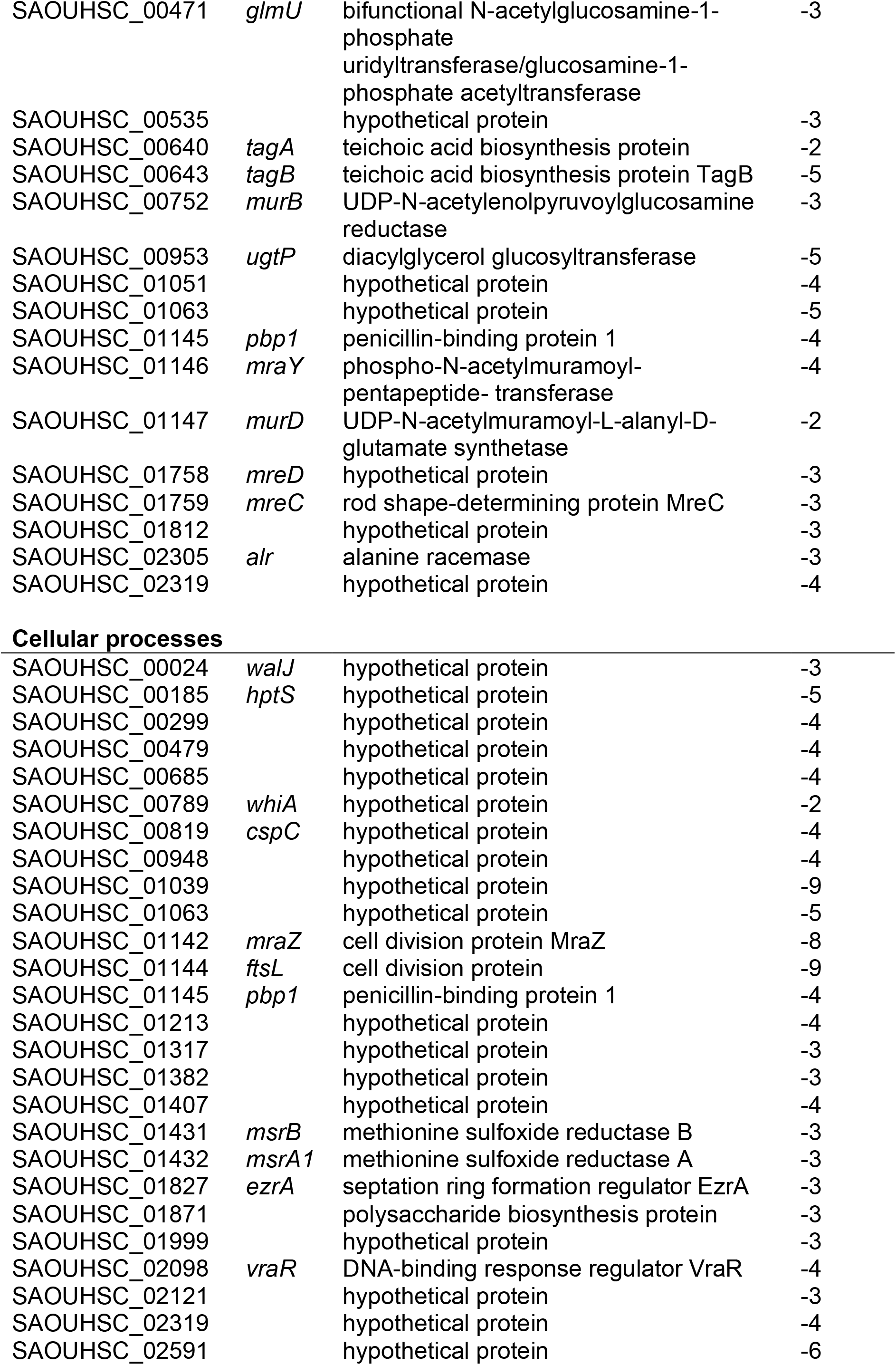
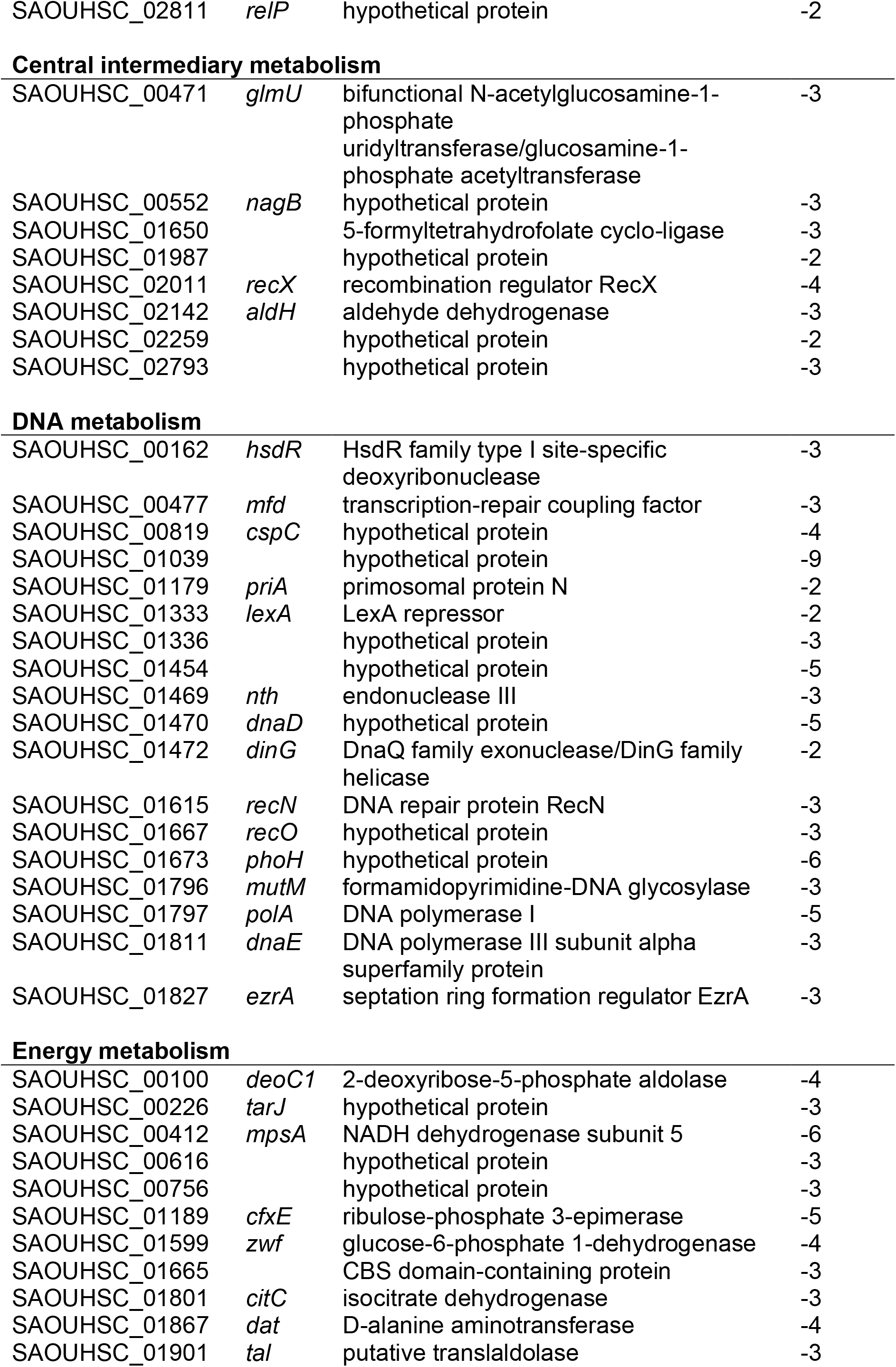
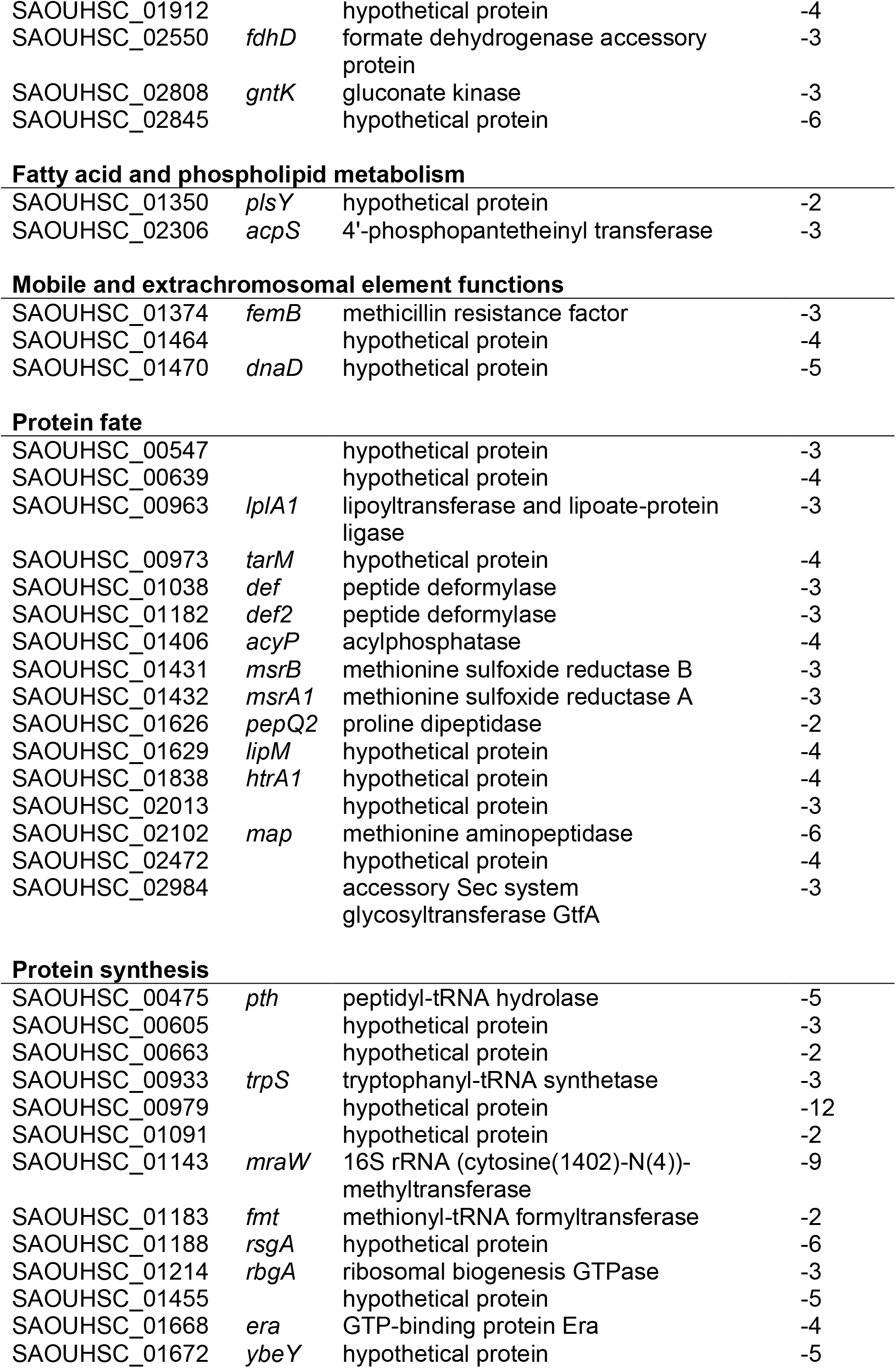
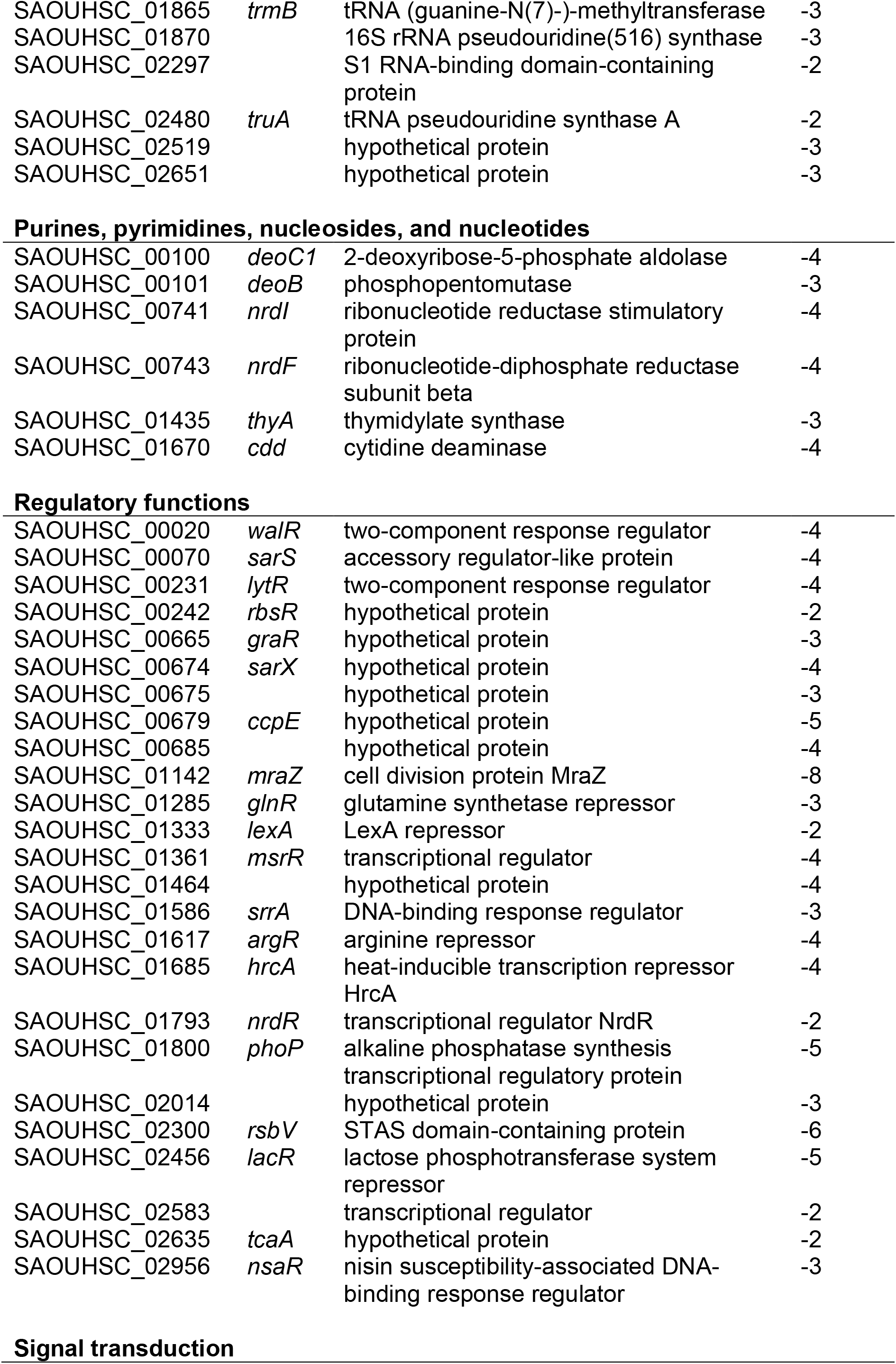
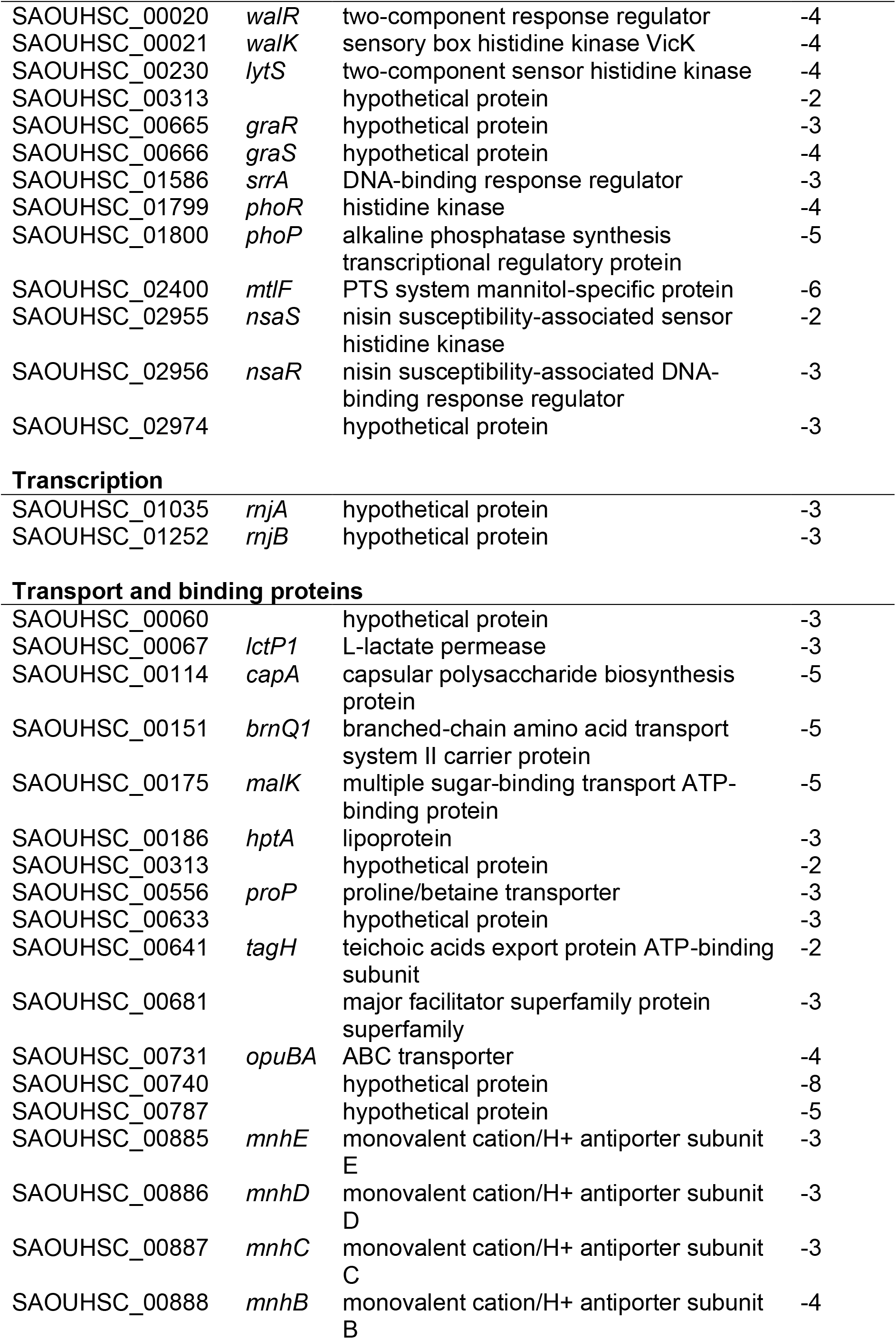
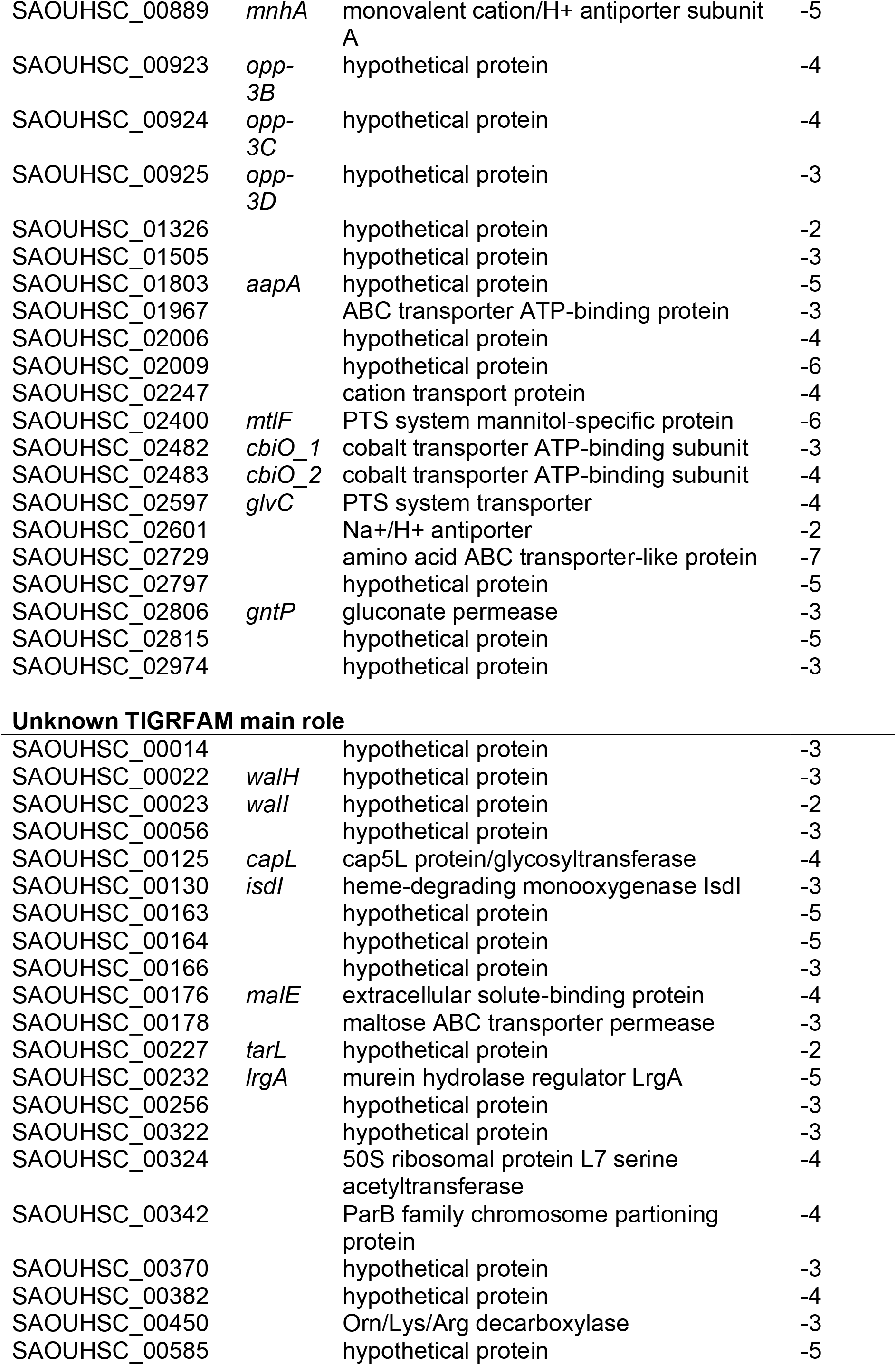
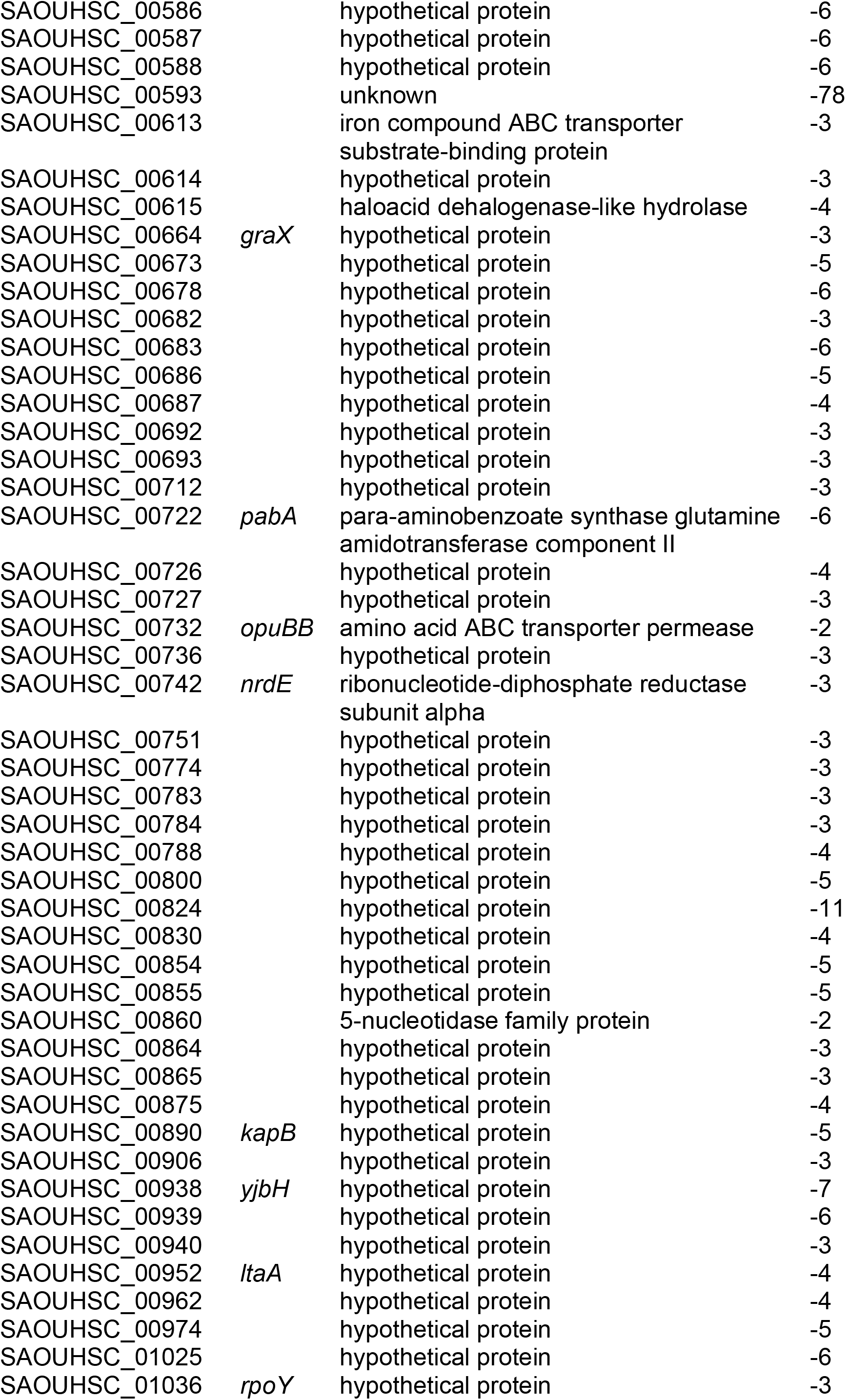
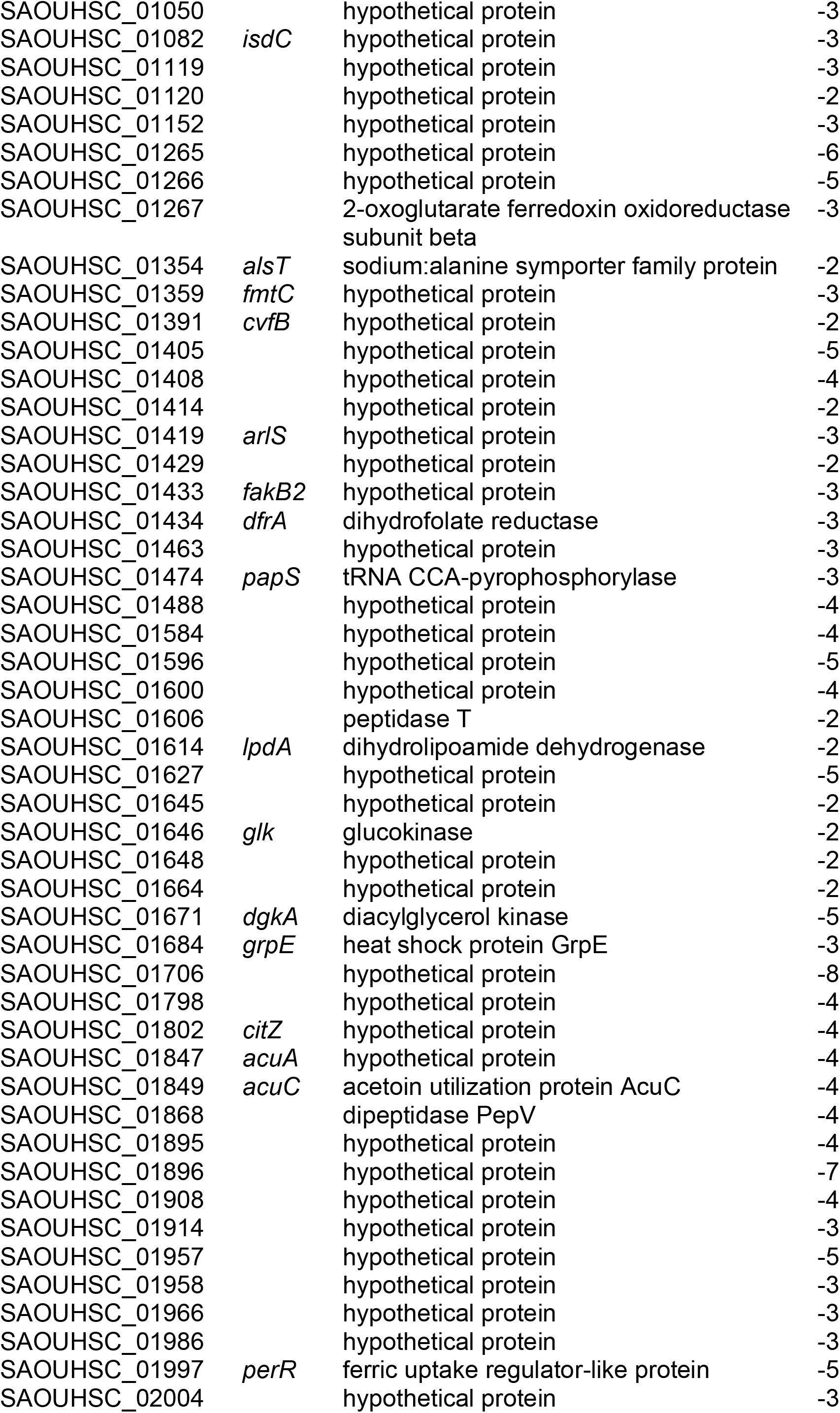
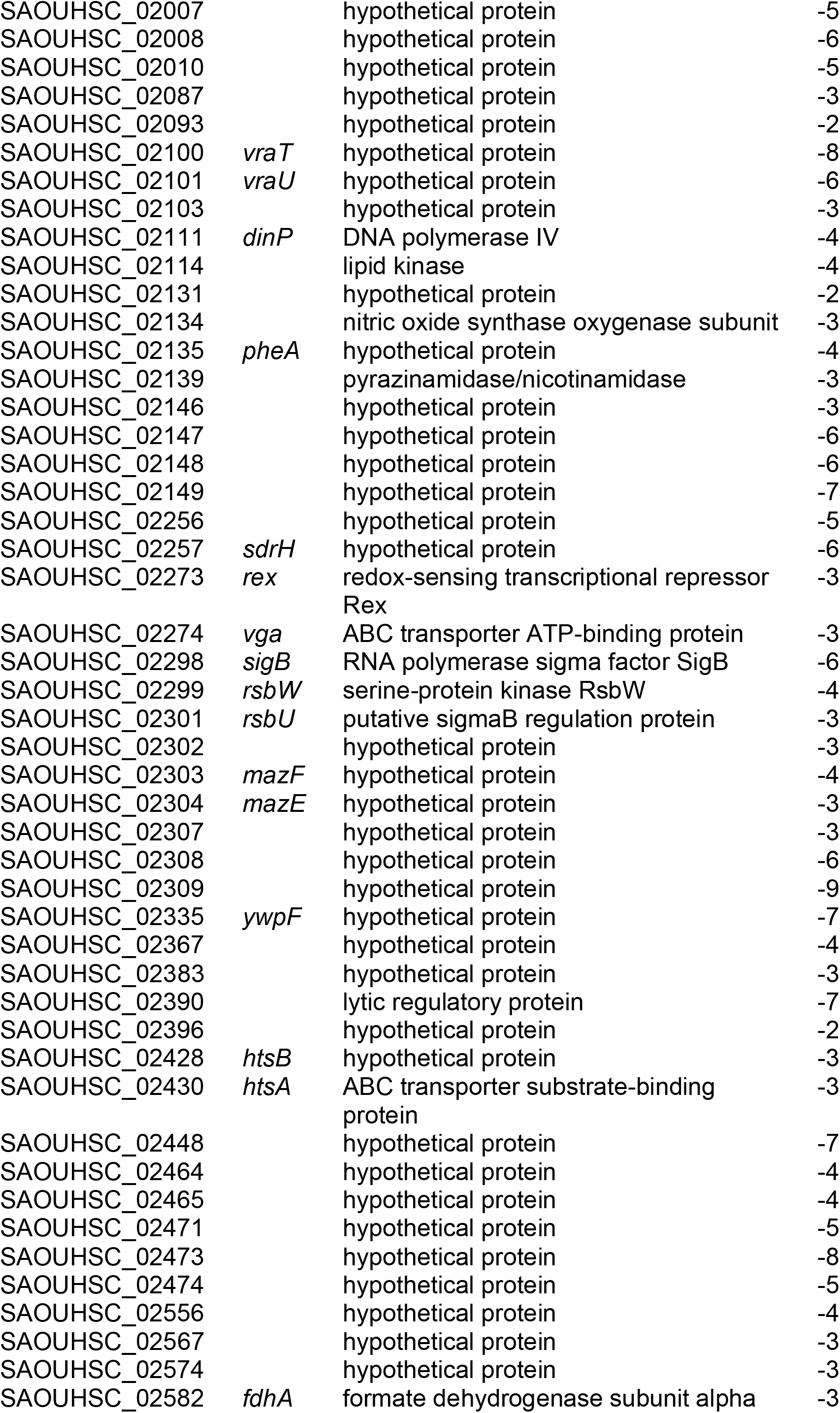
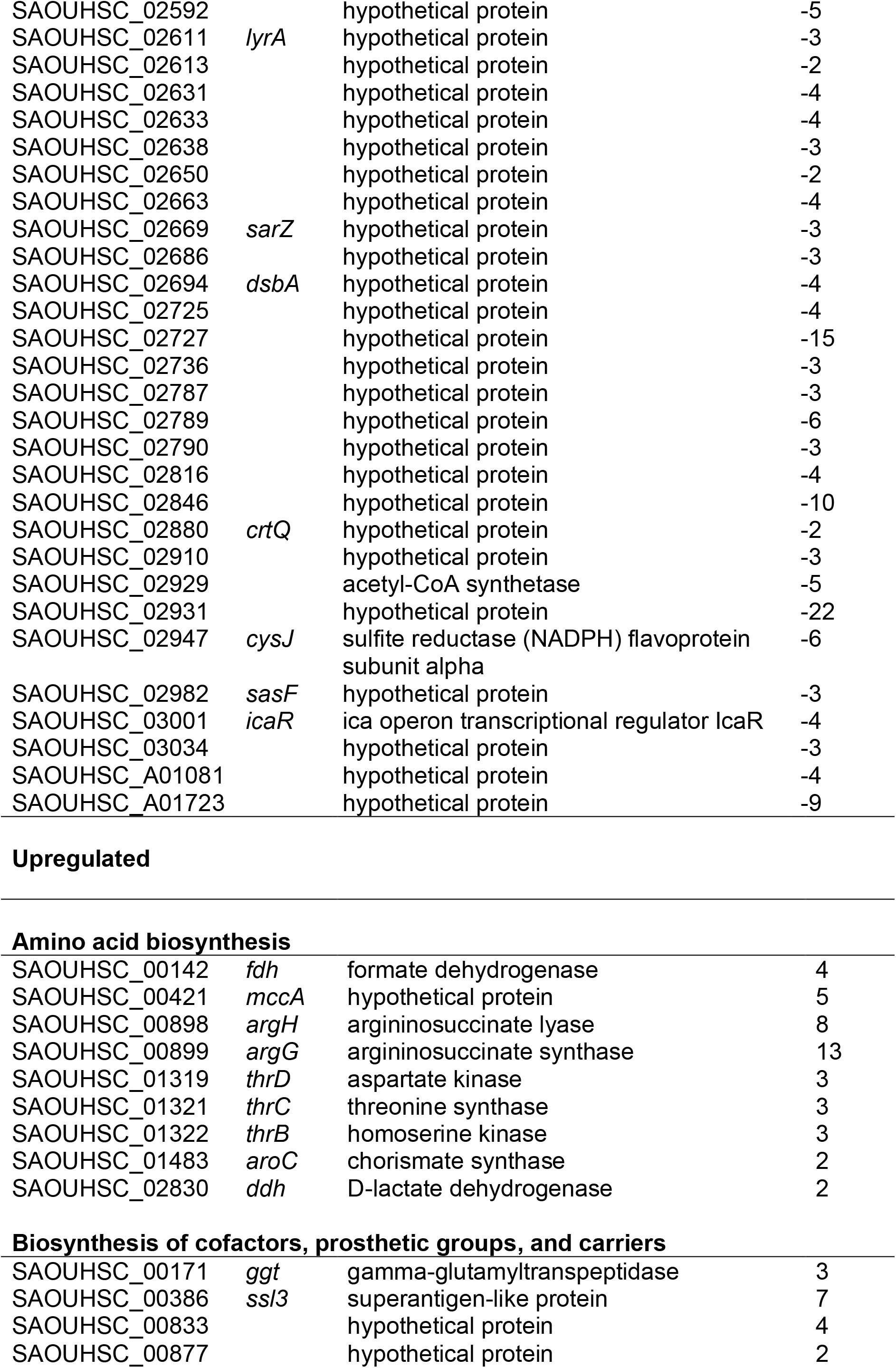
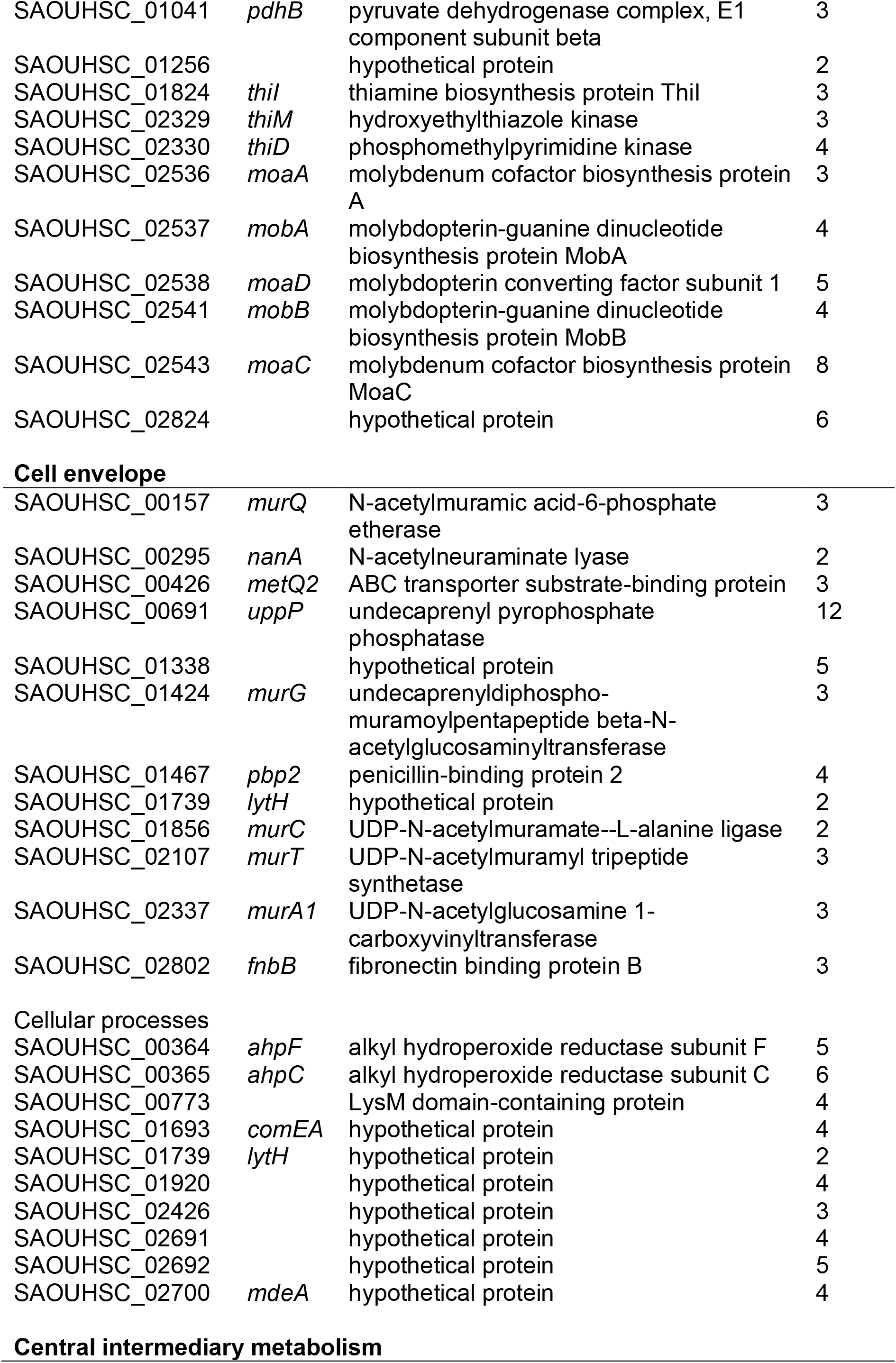
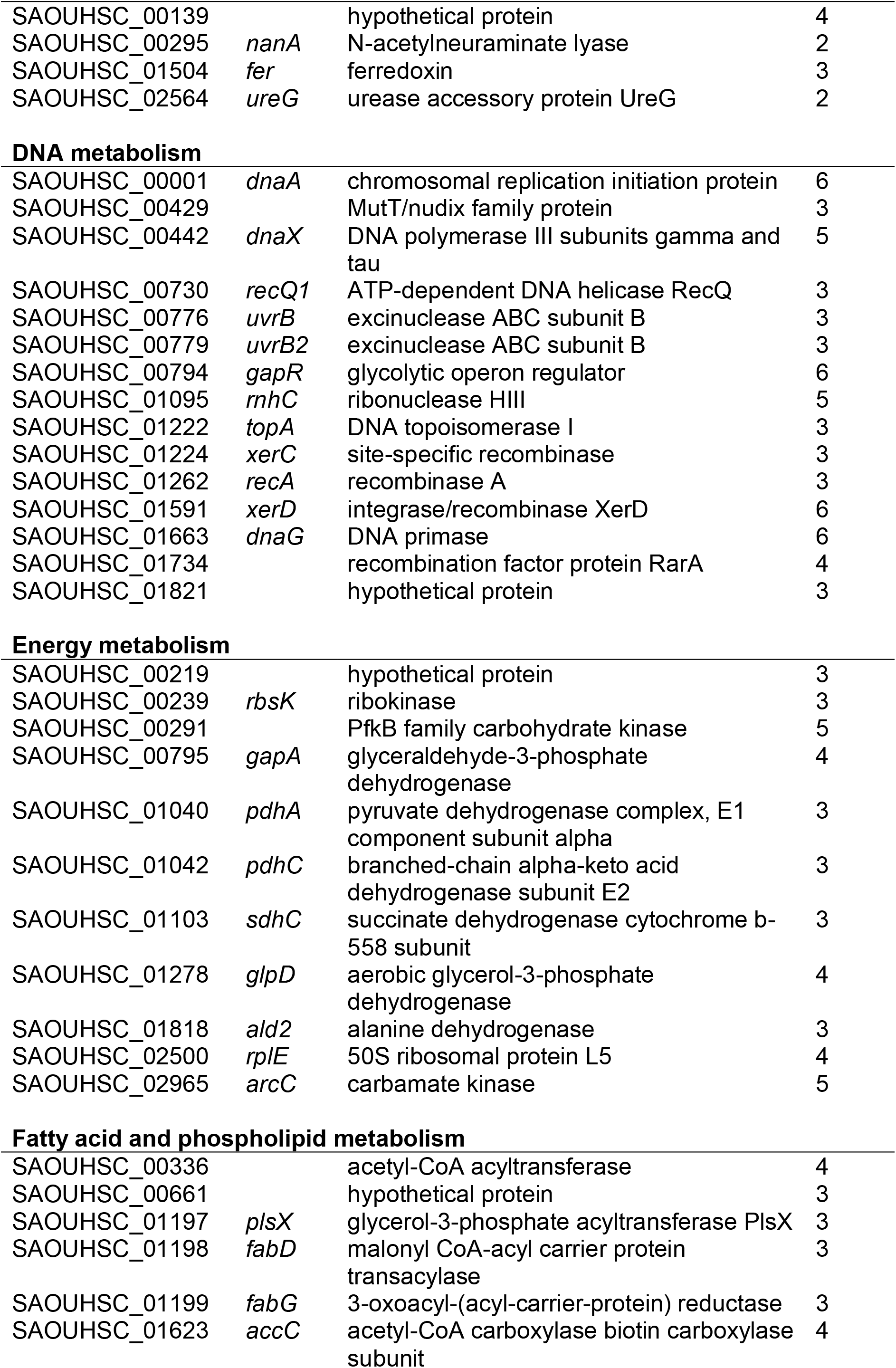
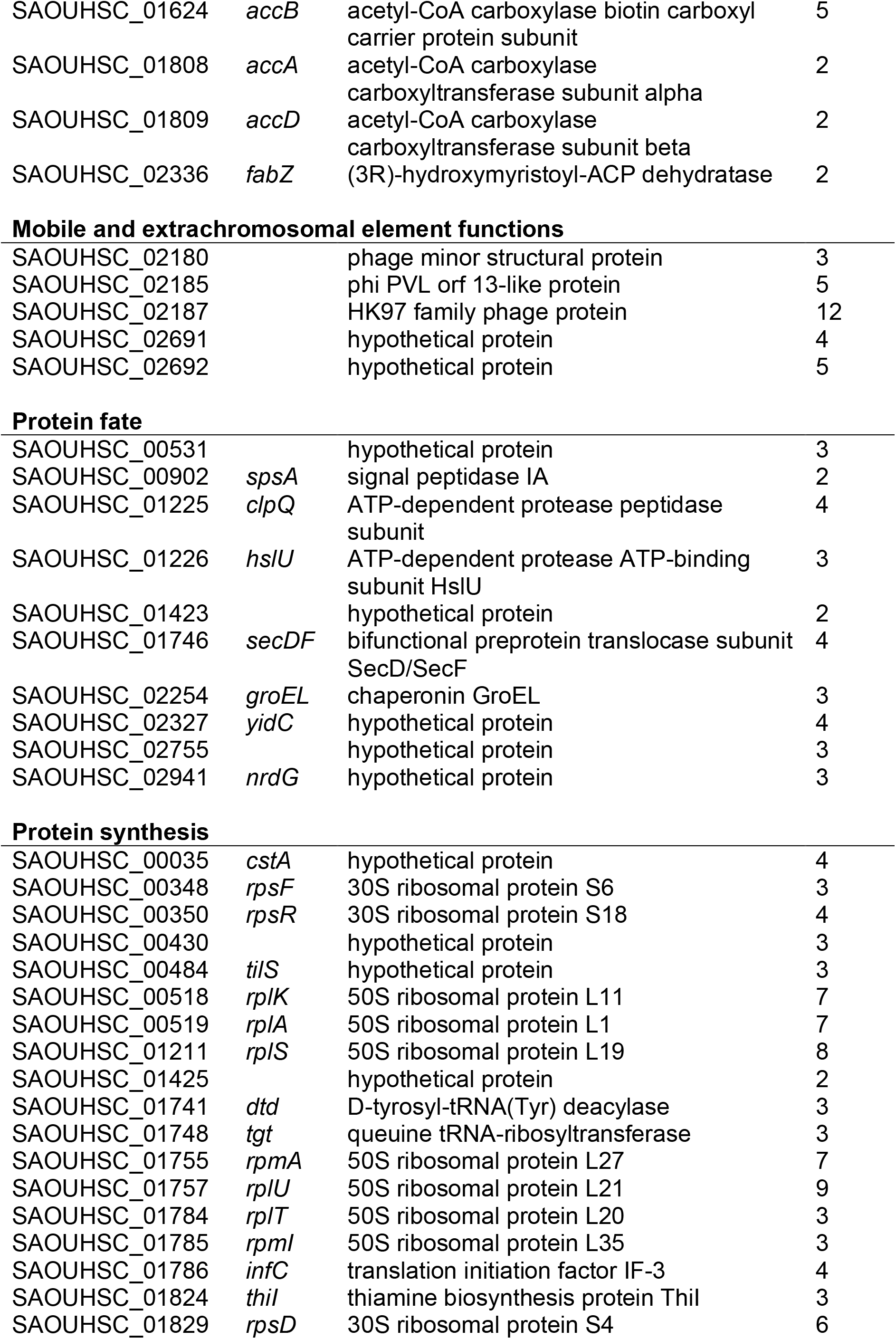
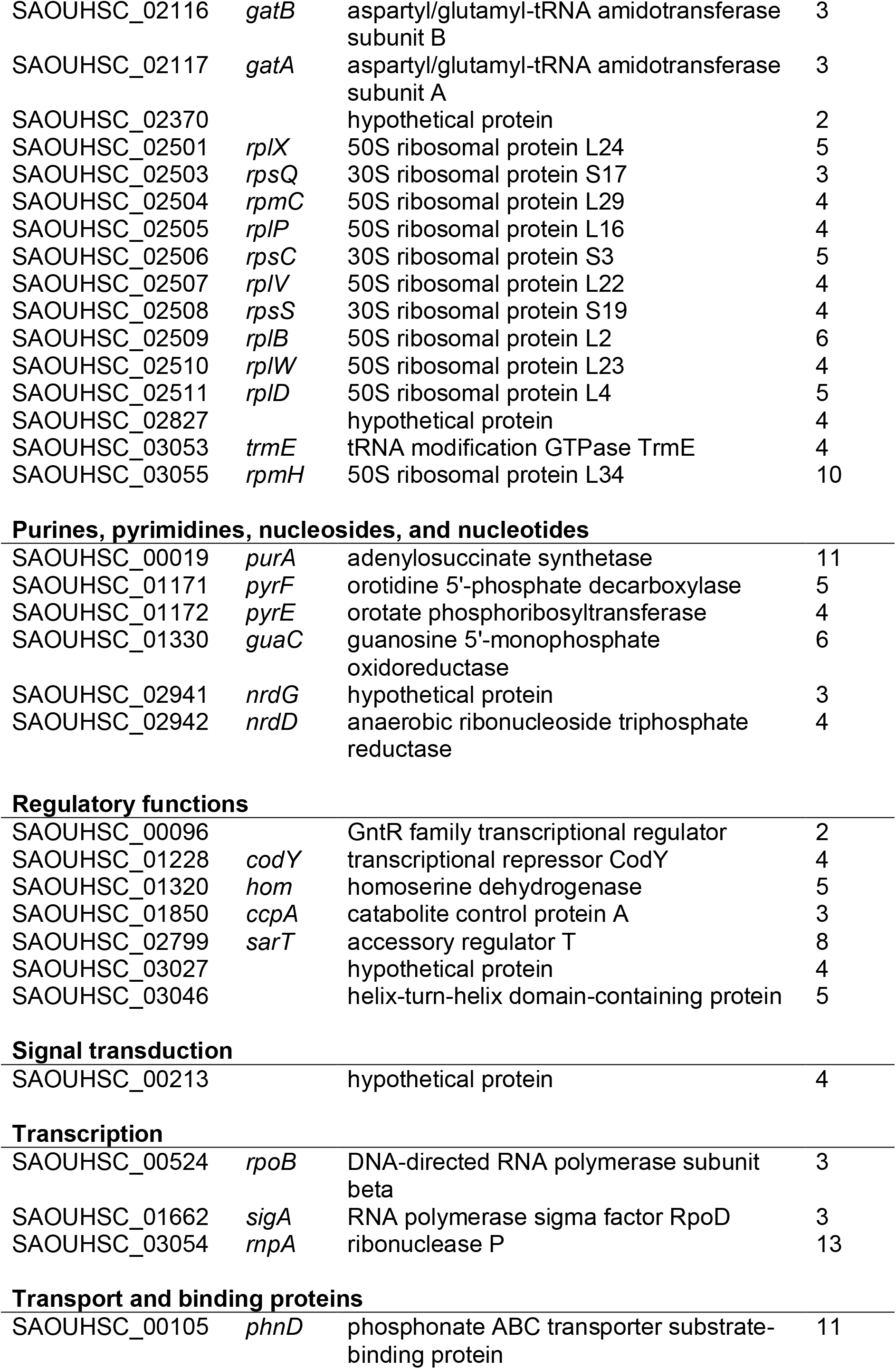
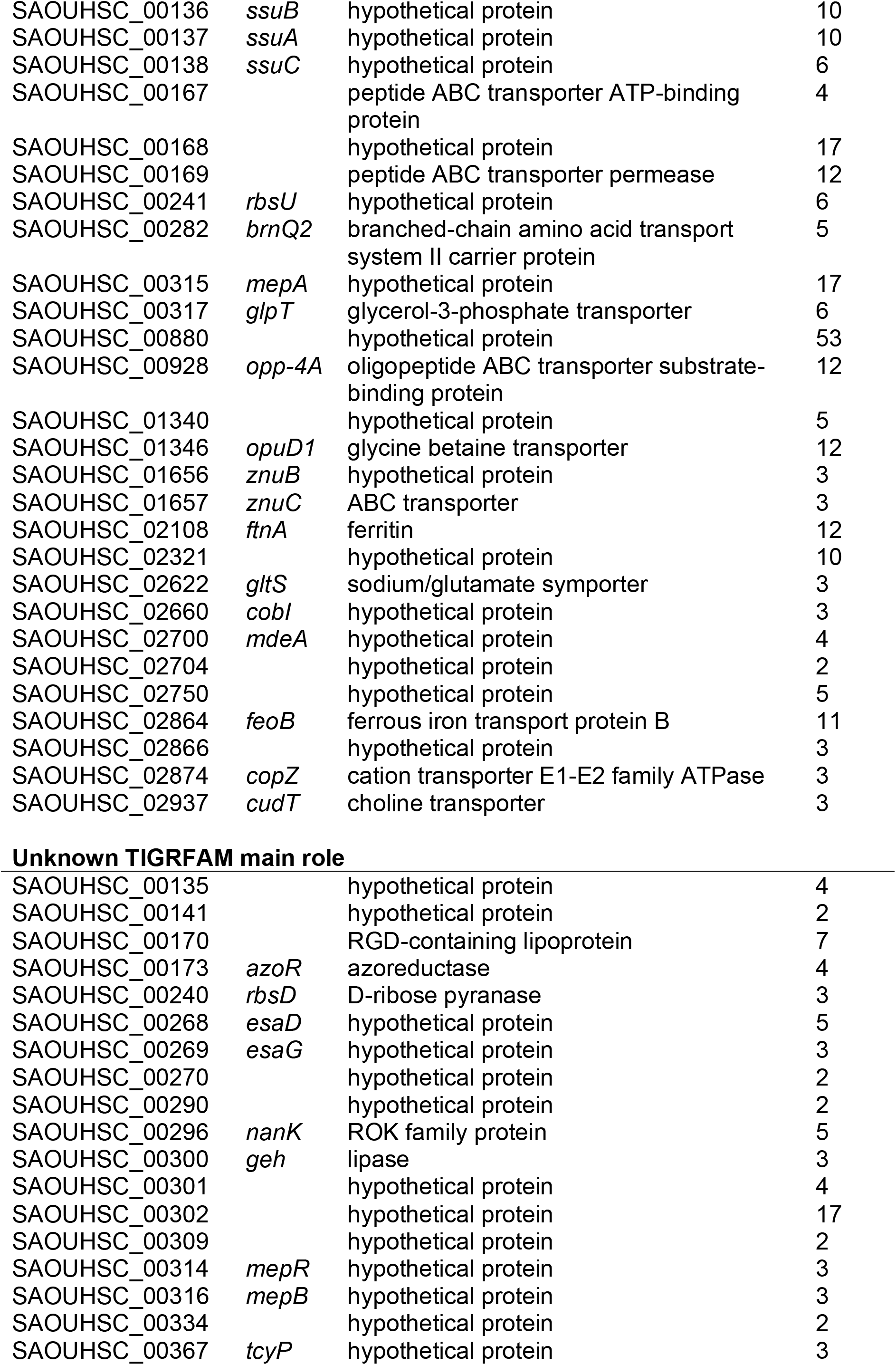
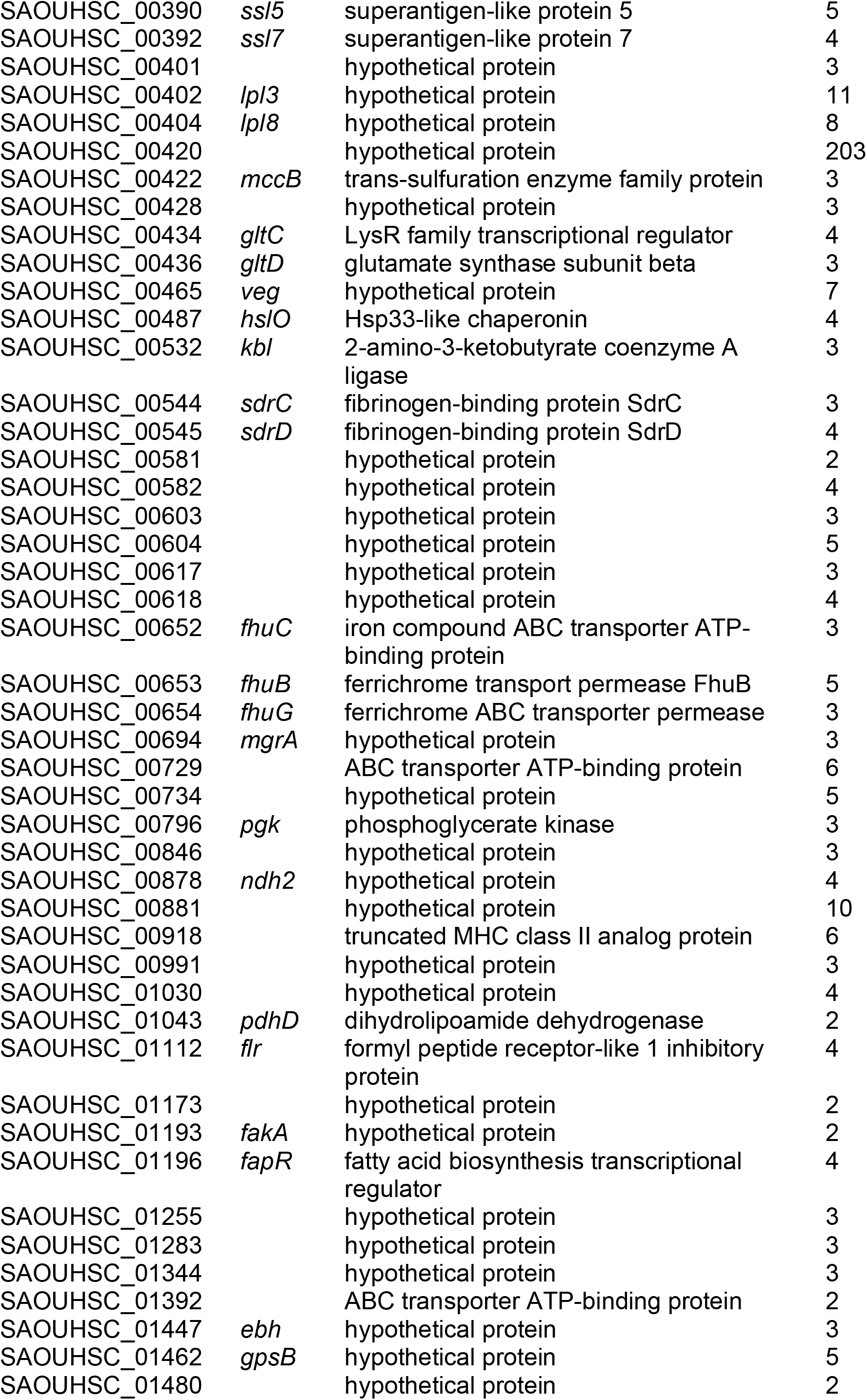
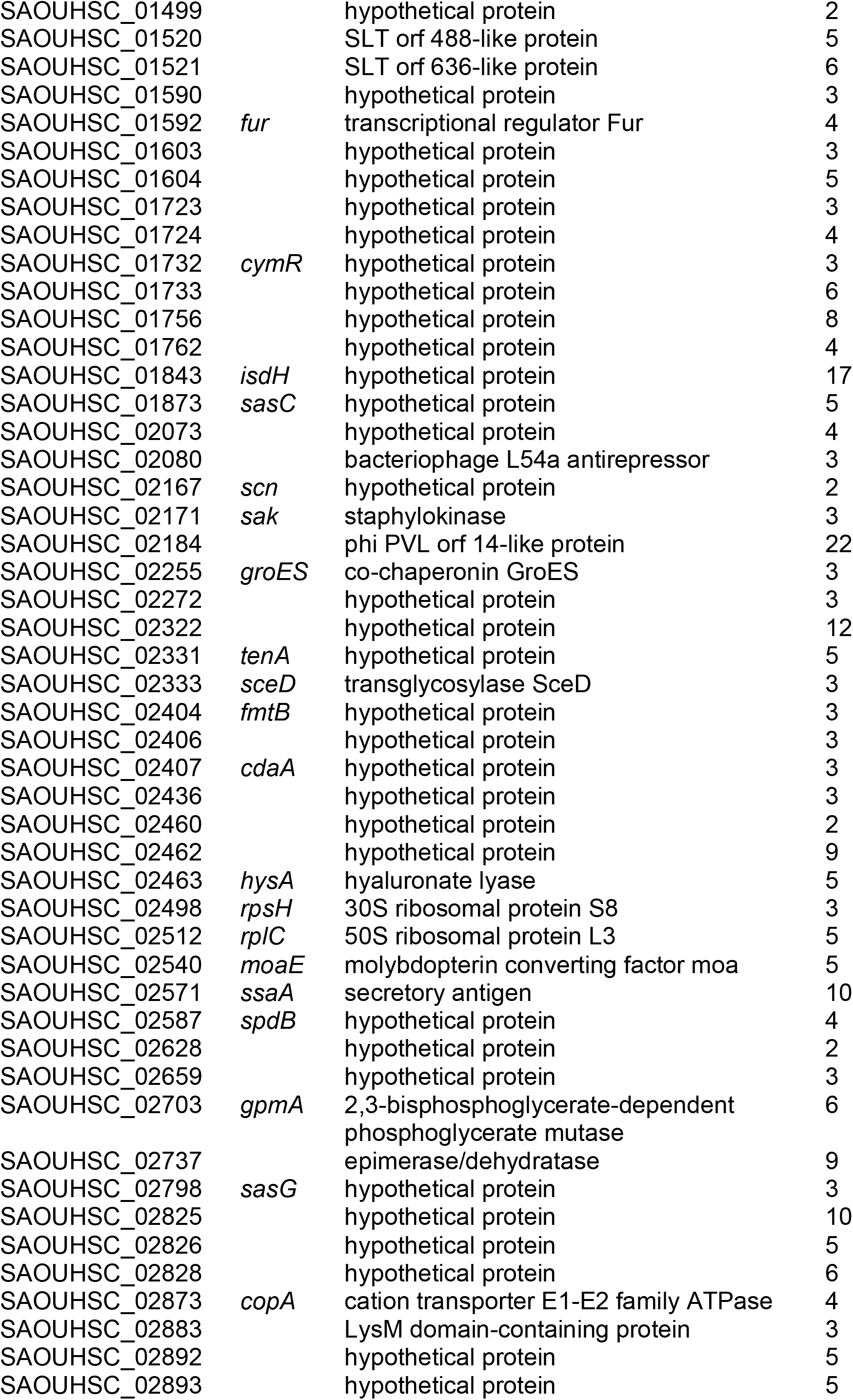
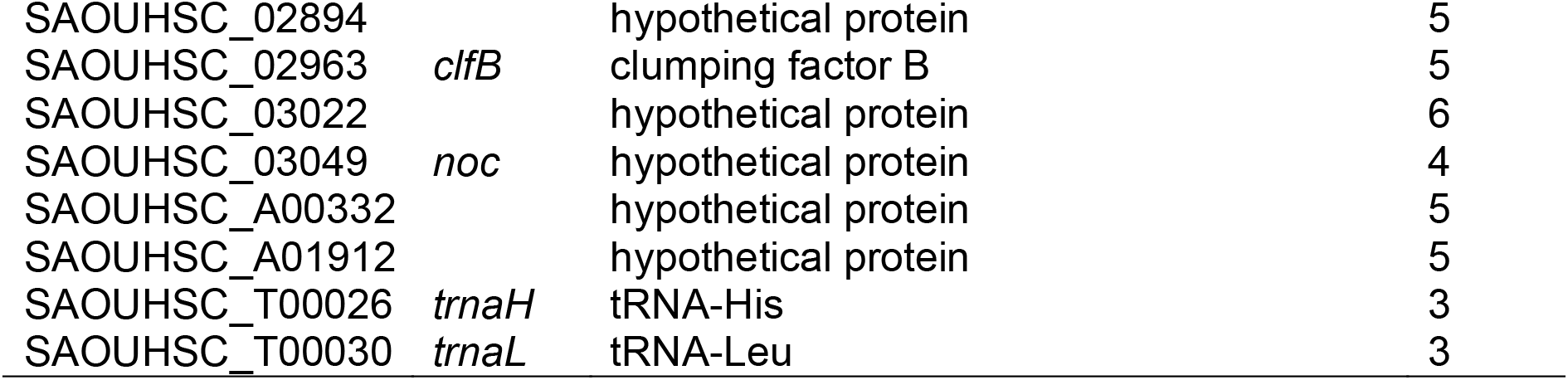
List of genes differentially expressed by a factor of ≥ 2 or ≤ −2 in MGB-BP-3 treated *S. aureus NCTC8325* compared to non-treated controls after 10 min of exposure.

## References

1. Barrett, M. P., Gemmell, C. G. & Suckling, C. J. Minor groove binders as anti-infective agents. Pharmacol. Ther. 139, 12–23 (2013).

2. Ross, W., Ernst, A. & Gourse, R. L. Fine structure of E. coli RNA polymerase-promoter interactions: alpha subunit binding to the UP element minor groove. Genes Dev. 15, 491–506 (2001).

3. Cai, X., Gray, P. J., Jr. & Von Hoff, D. D. DNA minor groove binders: back in the groove. Cancer Treat. Rev. 35, 437–450 (2009).

4. Puschendorf, B. et al. Studies on the effect of distamycin A on the DNA dependent RNA polymerase system. Biochem. Biophys. Res. Commun. 43, 617–624 (1971).

5. Taylor, A., Webster, K. A., Gustafson, T. A. & Kedes, L. The anti-cancer agent distamycin A displaces essential transcription factors and selectively inhibits myogenic differentiation. Mol. Cell. Biochem. 169, 61–72 (1997).

6. Anthony, N. G. et al. Antimicrobial lexitropsins containing amide, amidine, and alkene linking groups. J. Med. Chem. 50, 6116–6125 (2007).

7. Hlaka, L. et al. Evaluation of minor groove binders (MGBs) as novel anti-mycobacterial agents and the effect of using non-ionic surfactant vesicles as a delivery system to improve their efficacy. J. Antimicrob. Chemother. 72, 3334–3341 (2017).

8. Scott, F. J. et al. Selective anti-malarial minor groove binders. Bioorg. Med. Chem. Lett. 26, 3326–3329 (2016).

9. Scott, F. J. et al. An evaluation of Minor Groove Binders as anti-Trypanosoma brucei brucei therapeutics. Eur. J. Med. Chem. 116, 116–125 (2016).

10. Scott, F. J. et al. An evaluation of Minor Groove Binders as anti-fungal and anti-mycobacterial therapeutics. Eur. J. Med. Chem. 136, 561–572 (2017).

11. Scott, F. J. et al. An evaluation of Minor Groove Binders as anti-lung cancer therapeutics. Bioorg. Med. Chem. Lett. 26, 3478–3486 (2016).

12. Anthony, N. G. et al. Short lexitropsin that recognizes the DNA minor groove at 5’-ACTAGT-3’: understanding the role of isopropyl-thiazole. J. Am. Chem. Soc. 126, 11338–11349 (2004).

13. Koressaar, T. & Remm, M. Enhancements and modifications of primer design program Primer3. Bioinformatics 23, 1289–1291 (2007).

14. Untergasser, A. et al. Primer3--new capabilities and interfaces. Nucleic Acids Res. 40, e115 (2012).

15. Kolb, A., Kotlarz, D., Kusano, S. & Ishihama, A. Selectivity of the Escherichia coli RNA polymerase E sigma 38 for overlapping promoters and ability to support CRP activation. Nucleic Acids Res. 23, 819–826 (1995).

16. Rossiter, A. E. et al. Expression of different bacterial cytotoxins is controlled by two global transcription factors, CRP and Fis, that co-operate in a shared-recruitment mechanism. Biochem. J. 466, 323–335 (2015).

17. Browning, D. F. et al. The Escherichia coli K-12 NarL and NarP proteins insulate the nrf promoter from the effects of integration host factor. J. Bacteriol. 188, 7449–7456 (2006).

18. Squire, D. J. et al. Competition between NarL-dependent activation and Fis-dependent repression controls expression from the Escherichia coli yeaR and ogt promoters. Biochem. J. 420, 249–257 (2009).

19. Reynolds, J. & Wigneshweraraj, S. Molecular insights into the control of transcription initiation at the Staphylococcus aureus agr operon. J. Mol. Biol. 412, 862–881 (2011).

20. Chaudhuri, R. R. et al. Comprehensive identification of essential Staphylococcus aureus genes using Transposon-Mediated Differential Hybridisation (TMDH). BMC Genomics 10, 291 (2009).

21. Fuchs, S. et al. AureoWiki The repository of the Staphylococcus aureus research and annotation community. Int. J. Med. Microbiol. 308, 558–568 (2018).

22. Smits, W. K., Merrikh, H., Bonilla, C. Y. & Grossman, A. D. Primosomal proteins DnaD and DnaB are recruited to chromosomal regions bound by DnaA in Bacillus subtilis. J. Bacteriol. 193, 640–648 (2011).

23. Huang, Y. H., Lien, Y., Huang, C. C. & Huang, C. Y. Characterization of Staphylococcus aureus Primosomal DnaD Protein: Highly Conserved C-Terminal Region Is Crucial for ssDNA and PriA Helicase Binding but Not for DnaA Protein-Binding and Self-Tetramerization. PLoS One 11, e0157593 (2016).

24. DeFrancesco, A. S. et al. Genome-wide screen for genes involved in eDNA release during biofilm formation by Staphylococcus aureus. Proc. Natl. Acad. Sci. U. S. A. 114, E5969–E5978 (2017).

25. Nuxoll, A. S. et al. CcpA regulates arginine biosynthesis in Staphylococcus aureus through repression of proline catabolism. PLoS Pathog. 8, e1003033 (2012).

26. Miller, C. M., Baumberg, S. & Stockley, P. G. Operator interactions by the Bacillus subtilis arginine repressor/activator, AhrC: novel positioning and DNA-mediated assembly of a transcriptional activator at catabolic sites. Mol. Microbiol. 26, 37–48 (1997).

27. Prados, J., Linder, P. & Redder, P. TSS-EMOTE, a refined protocol for a more complete and less biased global mapping of transcription start sites in bacterial pathogens. BMC Genomics 17, 849 (2016).

28. Choe, D. et al. Genome-scale analysis of Methicillin-resistant Staphylococcus aureus USA300 reveals a tradeoff between pathogenesis and drug resistance. Sci. Rep. 8, 2215 (2018).

29. Browning, D. F. & Busby, S. J. Local and global regulation of transcription initiation in bacteria. Nat. Rev. Microbiol. 14, 638–650 (2016).

30. Sasse-Dwight, S. & Gralla, J. D. Probing co-operative DNA-binding in vivo. The lac O1:O3 interaction. J. Mol. Biol. 202, 107–119 (1988).

31. Rahman, A., O’Sullivan, P. & Rozas, I. Recent developments in compounds acting in the DNA minor groove. MedChemComm 10, 26–40 (2019).

32. Theuretzbacher, U. et al. Analysis of the clinical antibacterial and antituberculosis pipeline. Lancet Infect. Dis. 19, e40–e50 (2019).

33. Coates, A. R., Halls, G. & Hu, Y. Novel classes of antibiotics or more of the same? Br. J. Pharmacol. 163, 184–194 (2011).

