## Supplementary data; Resistance for "Novel antibiotic mode of action by repression of promoter isomerisation"

### Slide 1
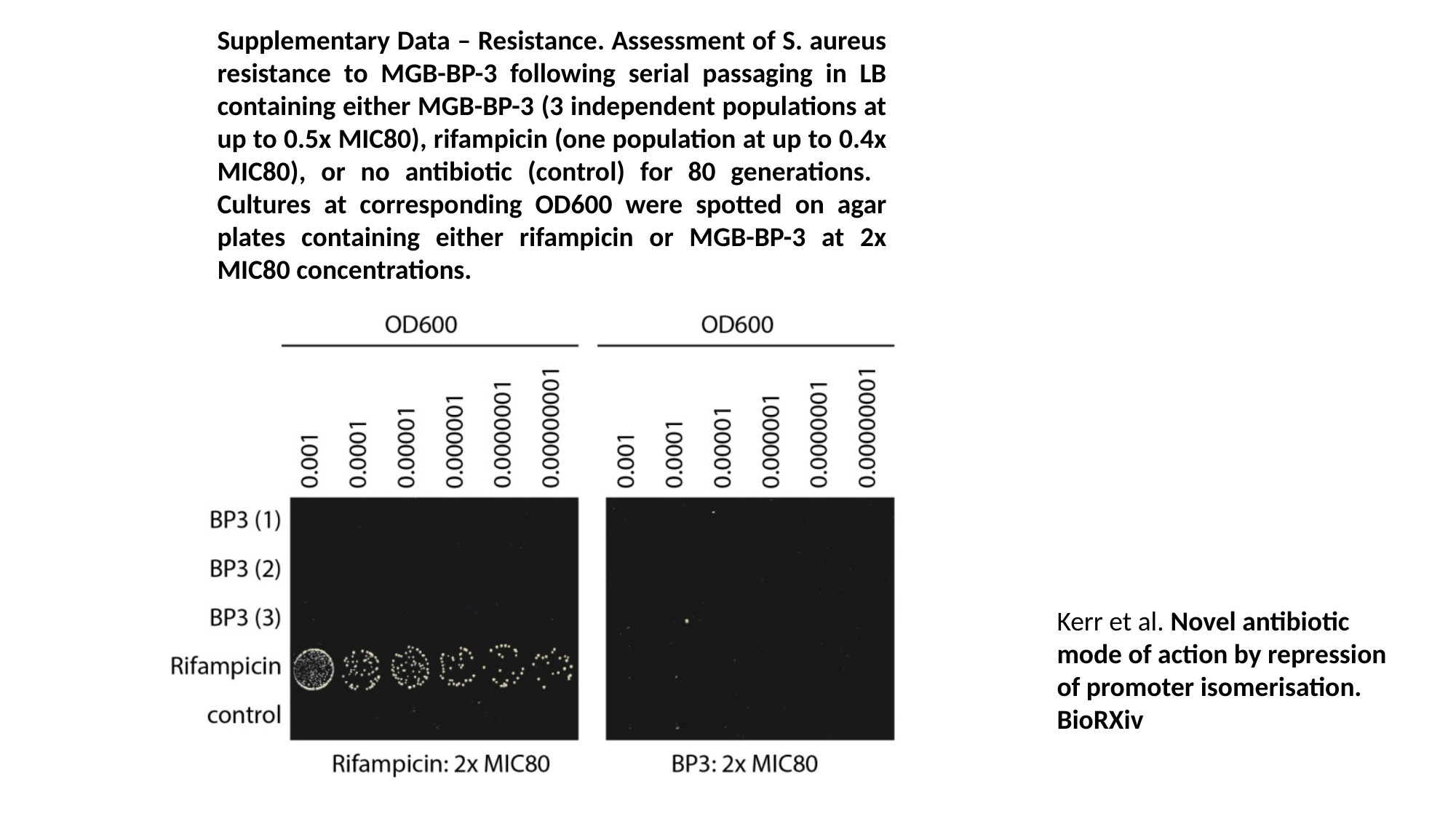

Supplementary Data – Resistance. Assessment of S. aureus resistance to MGB-BP-3 following serial passaging in LB containing either MGB-BP-3 (3 independent populations at up to 0.5x MIC80), rifampicin (one population at up to 0.4x MIC80), or no antibiotic (control) for 80 generations. Cultures at corresponding OD600 were spotted on agar plates containing either rifampicin or MGB-BP-3 at 2x MIC80 concentrations.
Kerr et al. Novel antibiotic mode of action by repression of promoter isomerisation.
BioRXiv
